# Phenylethynylbenzyl-Modified Biguanides Inhibit Pancreatic Cancer Tumor Growth

**DOI:** 10.1101/793109

**Authors:** Audrey Hébert, Maxime Parisotto, Marie-Camille Rowell, Alexandra Doré, Ana Fernandez, Guillaume Lefrancois, Paloma Kalegari, Gerardo Ferbeyre, Andreea R. Schmitzer

## Abstract

We present the design and synthesis of a small library of substituted biguanidium salts and their capacity to inhibit the growth/viability of pancreatic cancer cells. We first present their *in vitro* and membrane activity, before we address their mechanism of action in living cells and *in vivo* activity. We show that phenylethynyl biguanidium salts possess higher ability to cross hydrophobic barriers, improved mitochondrial accumulation and anticancer activity. Mechanistically, the most active compound **1b**, like metformin, decreases the NAD^+^/NADH ratio and mitochondrial respiration, but at 800-fold lower concentration. *In vivo* studies with, the most active, compound **1b** show a significant growth inhibition of pancreatic cancer xenografts in mice, while biguanides currently in clinical trials had no activity.

## Introduction

Although they have been used for decades in the treatment of type II diabetes, it is only quite recently that biguanides have been found to have interesting anticancer properties^1^. Many epidemiological studies have linked the long-term regular intake of metformin (1,1-dimethylbiguanide), a commonly used antidiabetic biguanide, to the reduction of the incidence of a variety of cancers in diabetic patients^2^. These results suggest that biguanides have anticancer activity, an idea that was experimentally verified using cancer models in cell culture and mice^3^. However, several clinical trials featuring metformin as a chemotherapeutic agent in humans have been unsuccessful ^4–6^. One of the reasons brought forward to explain this failure is the high hydrophilicity of metformin, administrated as a monoprotonated chloride salt at physiological pH (pKa 2.8 and 11.5). While animal models were given very high quantities of metformin to observe a potent anticancer activity^7^, the antidiabetic doses used in human patients were deemed too low to attain the reported antiproliferative concentration of 5 mM *in vitro*^8–9^.

Even if the molecular target of biguanides has not been identified yet, it is obvious that membrane insertion and mitochondrial penetration are critical for their activity. Different metformin analogues with lipophilic substituents and improved cellular penetration were previously reported. Narise *et al.* proposed in 2014 a series of phenformin derivatives possessing various substituents on the phenyl ring and bioisosteric replacements of the biguanide unit^10^. Neuzil *et al.* showed that biguanides functionalized with a mitochondria-targeting moiety such as triphenylphosphonium (TPP^+^), possess anticancer activities up to a 800-fold higher than metformin^11^. In agreement, it has been shown that metformin exert its anti-cancer activity at least in part by perturbing mitochondrial respiration ^7, 12^.

We have actively investigated the membrane perturbation properties of synthetic amphiphilic cationic ion transporters and antibiotics, including derivatives of phenylethynylbenzyl (PEB) – disubstituted imidazolium and benzimidazolium salts. We have demonstrated that the activities of these organic salts in artificial phospholipid bilayers or living prokaryotic and eukaryotic cells were the results of membrane penetration, self-assembly and partition^13^. Since all the studied organic salts containing a PEB unit in their structure were active on bacterial membranes and that mitochondria are essentially ancient bacteria trapped in eukaryotic cells, we reasoned that conjugating biguanides with the PEB unit could improve metformin’s cellular/mitochondrial uptake. We generated a small library of PEB-substituted biguanidium salts and their hydrogenated analogues and studied the capacity of these compounds to affect the growth/viability of pancreatic cancer cells. We identified a novel class of biguanide compounds with better membrane crossing abilities and more potent anticancer activity than metformin and phenformin.

## Results and Discussion

Phenylethynylbenzyl (PEB) – disubstituted imidazolium and benzimidazolium were shown to self-assemble inside phospholipid membranes and form stable channels through π-π interactions (Figure 1A)^13^. The transport of protons and ions through these open channels induced membrane perturbation, depolarization and bacterial death. Even if the PEB-disubstituted benzimidazolium salts were very active on bacterial membranes, their potential as mitochondrial membrane perturbators was not further explored because of their high toxicity on red blood cells (HC_50_ at 0.15 mM)^13–14^. The replacement of one PEB unit with a phenyl or a methyl group resulted in less toxic compounds (HC_50_ at 1 mM for the benzyl and 150 mM for the methyl substituent), forming more compact aggregates in the membranes.

**Figure 1:**
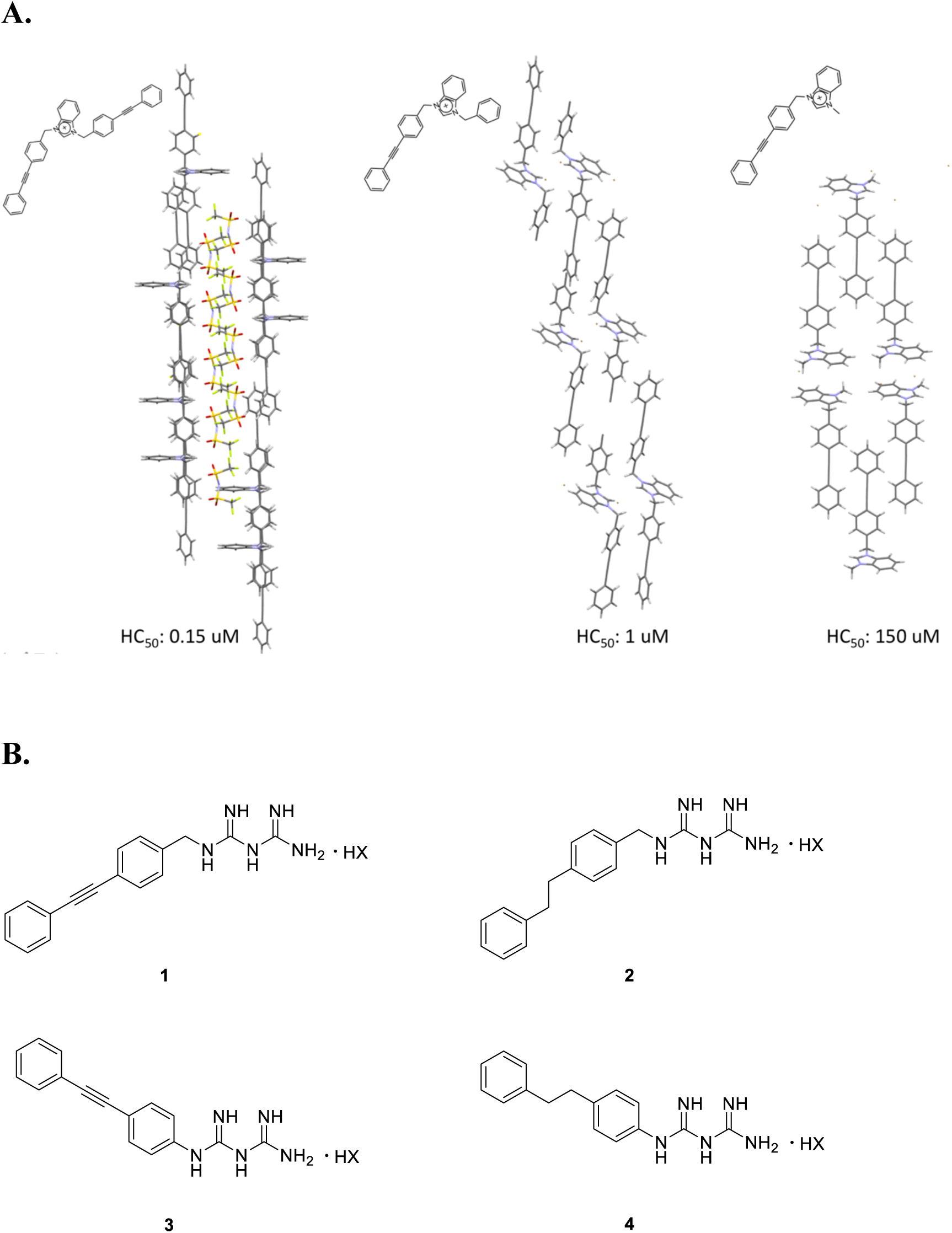
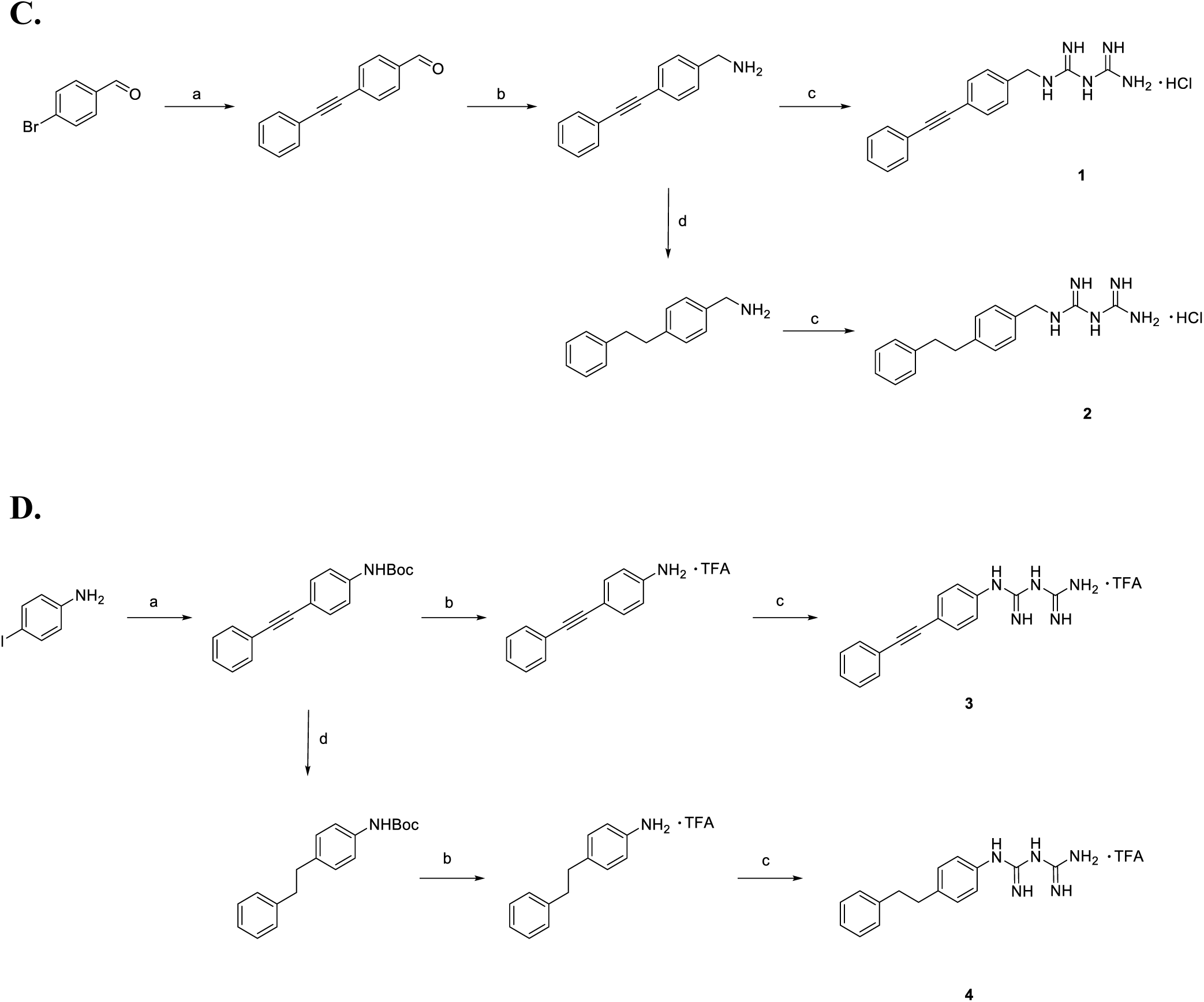
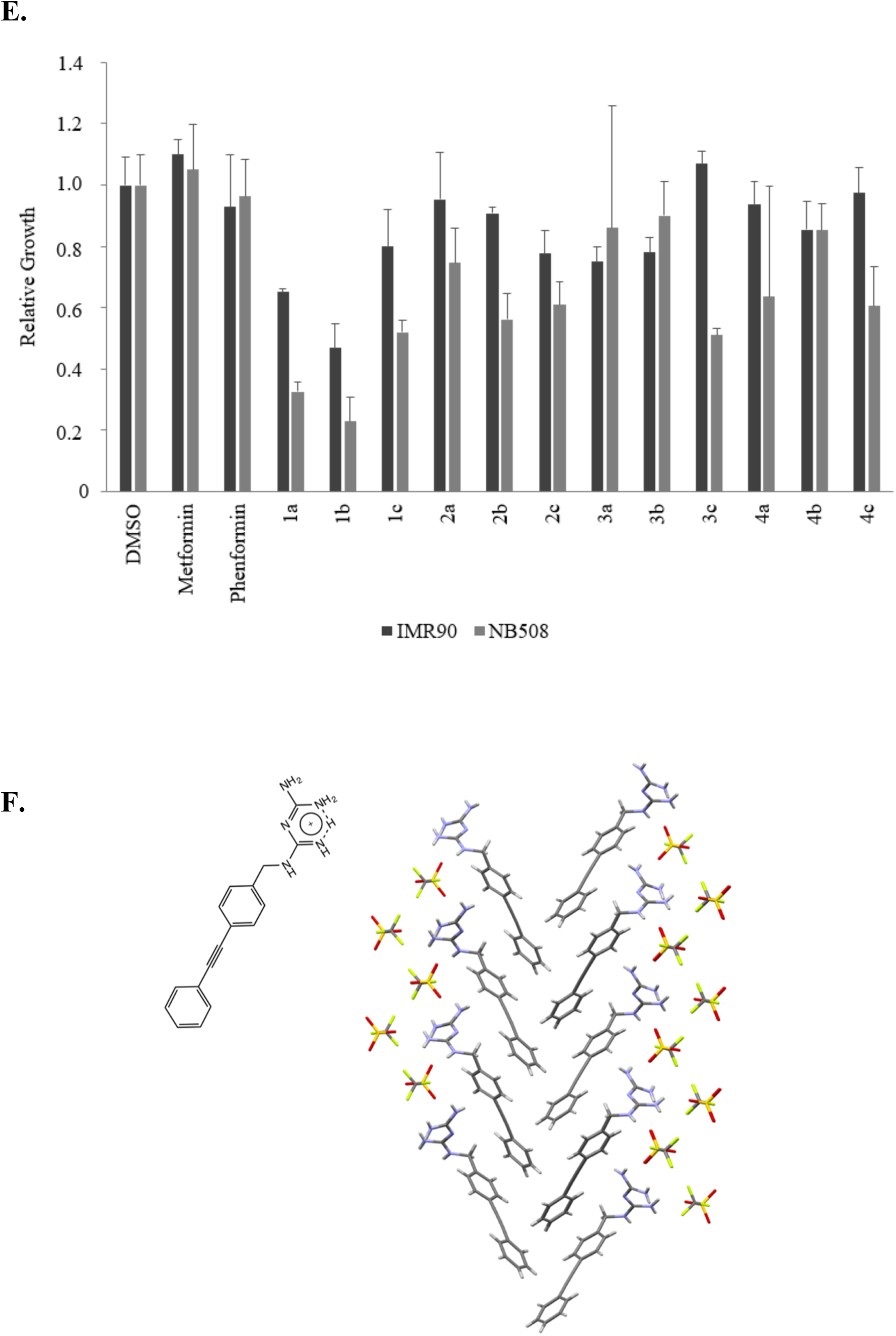
**A) Structure of previously studied PEB-substituted benzimidazolium salts^13^ B) Structure of PEB-substituted biguanidium salts. C) Synthesis of 4-(phenylethynyl)benzylbiguanide 1 and 4-(phenylethy)benzylbiguanide 2** (a) Phenylacetylene, PdCl_2_PPH_3_, CuI, PPH_3_, Et_3_N, THF, 70°C, o.n. (b) NaCN(BH_3_)_3_, EtOH_NH4OAc sat._/NH_3_, 80°C, o.n (c) Dicyandiamide, TMSCl, THF_anh._, 145°C, 1h (d) Pd/C 10%, H_2_, EtOH/AcOEt, 60°C, 2h. **D) Synthesis of 4-(phenylethynylphenyl)biguanidium 3 and 4-(phenylethylphenyl)biguanidium 4** (a) 1) Boc_2_O, THF, 2h, 2) Phenylacetylene, PdCl_2_PPH_3_, CuI, PPH_3_, Et_3_N, THF, 70°, o.n. (b) TFA, DCM, 60°C, 2h (c) Dicyandiamide, TMSCl, THF_anh._, 145°C, 1h (d) Pd/C 10%, H_2_, EtOH/AcOEt, 60°C, 2h. **E) Relative growth of NB508 mouse pancreatic ductal adenocarcinoma cells and IMR90 fibroblasts** exposed to 5 µM of PEB-biguanidium. Cells were incubated for 72 h at 37°C. **F) Crystal structure of 1b showing its self-assembly in the solid state**.

### Synthesis

PEB-Biguanidium chloride **1c** was synthesised by the cross-coupling of 4-bromobenzaldehyde with phenylacetylene followed by the reductive amination of the aldehyde with NaCNBH_3_. The biguanide was then formed by reacting the amine with dicyandiamide to afford PEB-biguanidium chloride **1c** with 30% yield. PEB-biguanidium chloride **2c** was obtained by hydrogenation of 4-(phenylethynyl)benzyl amine followed by the formation of the biguanidium chloride with 20% yield (Figure 1C).

PEB-biguanidium **3** was synthesized by the cross-coupling of BOC protected 4-iodoaniline with phenylacetylene followed by the deprotection of the amine and the formation of the biguanidium trifluoroacetate with dicyandiamide in a 24% yield. PEB-biguanidium **4** was synthesized by the hydrogenation of *tert*-butyl-(4-(phenylethynyl)phenyl)carbamate, deprotection of the amine and formation of the biguanidium trifluoroacetate with a 35% yield (Figure 1D).

The counter-anions of PEB-Biguanidium chloride **1c** and **2c** were exchanged through the anion metastasis of chloride with either LiNTf_2_ (**a**) or LiOTf (**b**) in methanol in quantitative yield. The counter-anions of PEB-biguanidium **3** and **4** were exchanged by the same procedure with the addition of a deprotonation step with NaHCO_3_ before the ion exchange step.

### *In vitro* screening

The biguanidium salts possessing various counter-anions were first screened for the growth inhibition of mouse pancreatic ductal adenocarcinoma cells NB508 and normal human fibroblasts IMR90 (Figure 1E), at 5 µM. Almost all the tested compounds were able to inhibit the growth of cancer cells at this concentration, while no effect was observed for metformin and phenformin at the same concentration. The most important antiproliferative activities were observed for **1a** and **1b**, both showing a good selectivity towards the cancer cells.

The toxicity of PEB-Biguanidiums salts was estimated through their hemolytic activity by incubating red blood cells (RBC) with the compounds for 1 hour (*See supporting information*). A very low hemolytic activity (under 10%) was observed for all the compounds, even at concentrations up to 150 µM. As RBC are mainly composed of a plasma membrane enveloping hemoglobin, the low hemolysis observed indicates that PEB-Biguanidium salts are not disrupting RBC’s membranes, compared to the previously reported benzimidazolium salts^15–16^.

### Membrane activity

As compounds **1a-c** showed good inhibition of the growth of cancer cells and low activity on fibroblasts, they were retained for further investigation. Compound **1b** was the only one with good methanol and DMSO solubility even at high concentrations, and was used for further investigation. The logP of compound **1b** (0.4) is much higher than metformin (−1.4)^17^ and phenformin (−0.8)^18^, indicating its higher hydrophobicity and higher membrane permeability capacity. The ability of **1b** to penetrate and cross phospholipid membranes, compared to metformin and phenformin, was studied by U-tube experiments, where two aqueous phases are separated by a bulk hydrophobic solvent such as chloroform, to mimick a bilayer membrane (*see supporting information*). A compound able to penetrate and cross a phospholipid bilayer is able in these conditions to partition into water and chloroform. A concentration of 250 µM of biguanide was added to the *cis*-aqueous side and the concentration of the *trans*-side was measured after 48h and 72h. Compound **1b** was able to partition rapidly into the chloroform and cross to the *trans*-aqueous side, while only traces of metformin and phenformin were measured even after 72h (Figure 2A).

**Figure 2:**
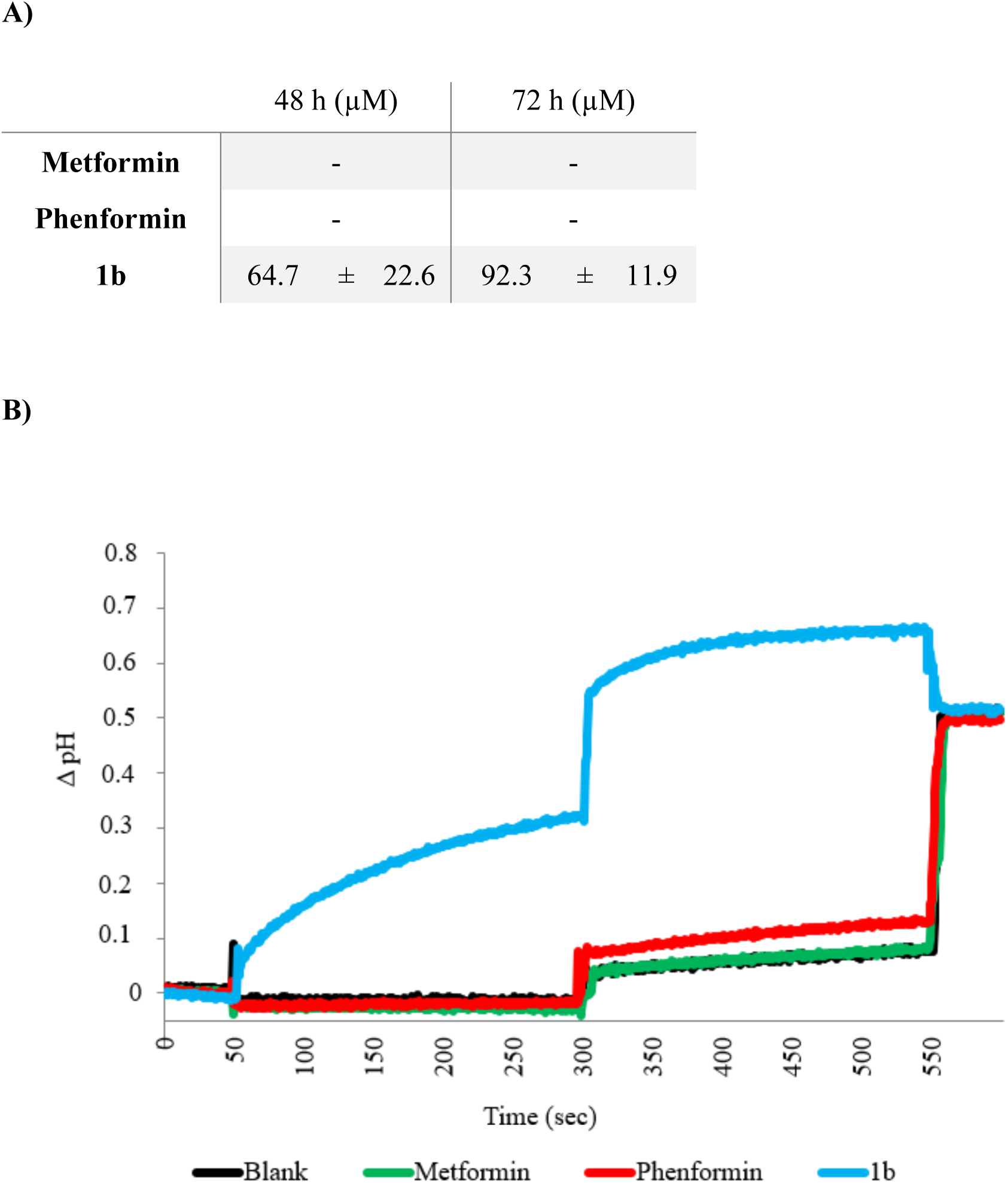
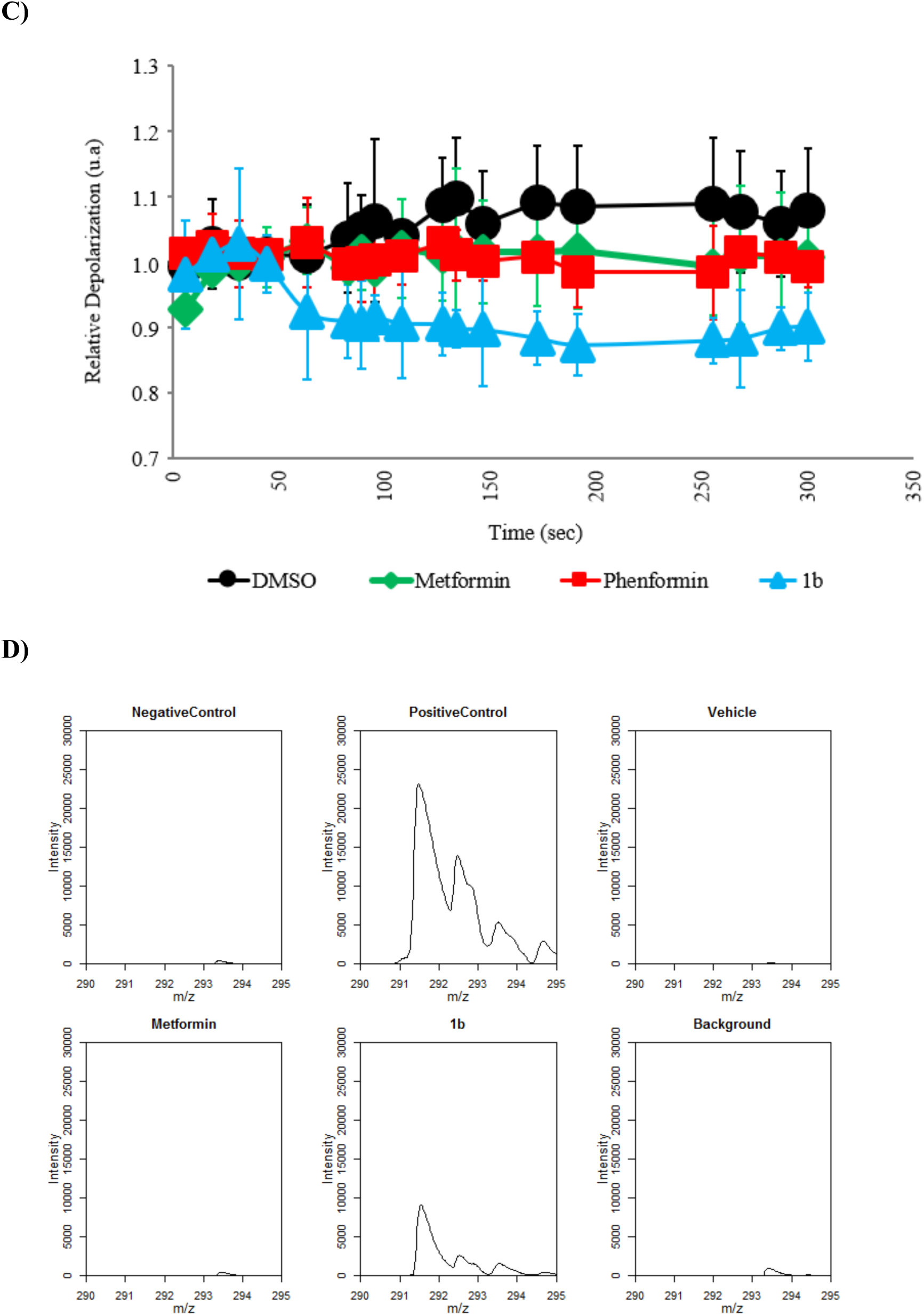
**A) Partition of biguanide in a U-tube experiment.** Concentration of biguanide on the *trans*-side of the U-tube at 48 h and 72 h at 25 °C, after addition of 250 µM of biguanide on the *cis*-side. **B) Variation in the internal pH of HPTS-containing EYPC liposomes.** Intravesicular solution: 1 mM HPTS, 10 mM HEPES, and 100 mM NaCl, adjusted to pH = 7.4, and extravesicular solution: 10 mM HEPES and 100 mM NaCl, adjusted to pH = 7.4. Biguanidiums were injected after 50 sec at 5 mM (50 mol% relative to the 10 mM EYPC concentration), a NaOH pulse was induced at 300 sec, and the liposomes were lysed with Triton-X at 550 sec. Each curve is the average of three independent measurements. **C) Fluorescence of safranin O in the EYPC liposomes.** Biguanidiums were injected at 5 mM (50 mol% relative to the 10 mM EYPC concentration) at 50 sec. Each curve is the average of three independent measurements. **D) Mitochondrial penetration of compound 1b and metformin.** Mitochondrial isolation was performed according to^19^. For each experiment, an anti-HA IP was performed on KP4 cells expressing pMXs-3XHA-EGFP-OMP25 that were treated for 3 hours with 15 μM metformin, 15 μM compound **1b** or vehicle.

Despite its structural similarity with the PEB-substituted benzimidazolium synthetic transporters and ability to penetrate phospholipid membranes, compound **1b** did not transport chloride across the phospholipid membrane of EYPC-LUVs (*See supporting information*). As it can be observed in the solid-state structure obtained from chloroform, **1b** does not form channel-like supramolecular architectures, but self-assembles into herringbone-shaped interlocked dimers that do not seem to possess chloride transport properties (Figure 1F).

However, when using EYPC-LUVs containing 8-Hydroxypyrene-1,3,6-trisulfonic acid trisodium salt (HPTS) as pH-sensitive probe, a change of the internal vesicular pH was observed when compound **1b** was added (Figure 2B). Initially, the pH was neutral and equal in both the extra and intra-vesicular solution. Upon injection of the compound we observed a basification of the internal liposomal solution. This basification was further enhanced with the addition of a base pulse to the external solution at 300 sec, indicating either a transport of protons to the extravesicular solution or the transport of hydroxyls to the intravesicular solution. This was also reported for alkylbiguanidium salts and was described as an electrogenic process, *i.e.* the transmembrane transport of an ion without the balance of the charge through the movement of another ion, which usually results in a charge transfer^15^. Compound **1b**, like alkylbiguanidium salts, was not able to transport chloride, so the electroneutral hypothesis of a H^+^/Cl^−^ antiport or OH^−^/Cl^−^ symport can be rejected and the hypothesis of the electrogenic transport mechanism can be suggested. Metformin and phenformin showed no membrane activity in these experiments.

The transport of H^+^/OH^−^ ions across the membrane is an electrogenic mechanism that usually causes the depolarization of the electrochemical gradient across phospholipid membranes. When EYPC liposomes containing intravesicular K^+^ ions are bathing in an extravesicular solution containing Na^+^ ions, a small membrane potential is generated at their membranes and the fluorescence of Safranin O can be used as a membrane potential probe^20^. The addition of **1b** to the external solution of these Safranin-O containing liposomes resulted in a decrease of the Safranin’s fluorescence intensity, indicating a depolarization of the membrane (Figure 2C).

Mitochondrial membrane potential together with the proton gradient are responsible for ATP production^21^ and maintenance of cellular health and function. Compounds able to penetrate mitochondria and alter their membrane potential are usually used as chemotherapeutics. Compound **1b**, but not metformin, was able to efficiently penetrate mitochondria after a short incubation of cells with 15 µM concentration of each drug (Figure 2D). This indicates that compound **1b** easily diffuses into mitochondria unlike metformin that requires a specialized transporter^22^. These experiments altogether indicate that **1b** is able to penetrate phospholipid and mitochondrial membranes and alter their potential.

### Antiproliferative activity *in* vitro and mechanism of action

In order to compare the antiproliferative/anticancer activity of compound **1b** to that of the most commonly studied biguanides derivatives metformin and pheformin, we performed growth assays with human PDAC cell lines over 3 days in the presence of various concentrations of these compounds. The results showed that both in KP4 cells (Figure 3A) and in Panc1 cells (Figure 3B), metformin has an IC_50_ of about 5 mM. Phenformin, with an hydrophobic phenyl group, has as lower IC_50_: 51 µM in KP4 and 180 µM in Panc1 cells. Compound **1b** showed much lower IC_50_ of 6.1 µM in KP4 cells and 15 µM in Panc1 cells, 100-800 fold lower than metformin and 8-12 fold lower than phenformin.

**Figure 3.**
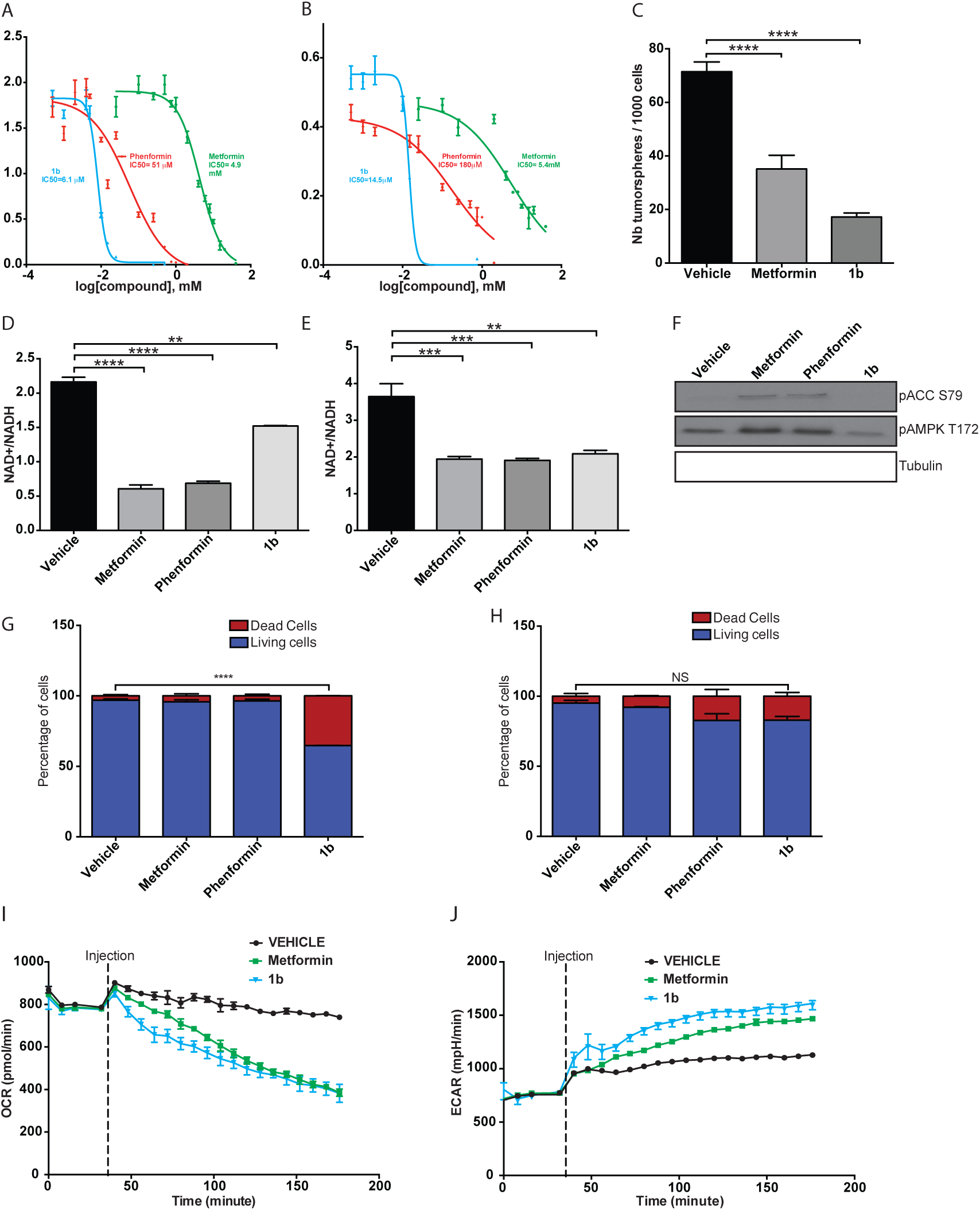
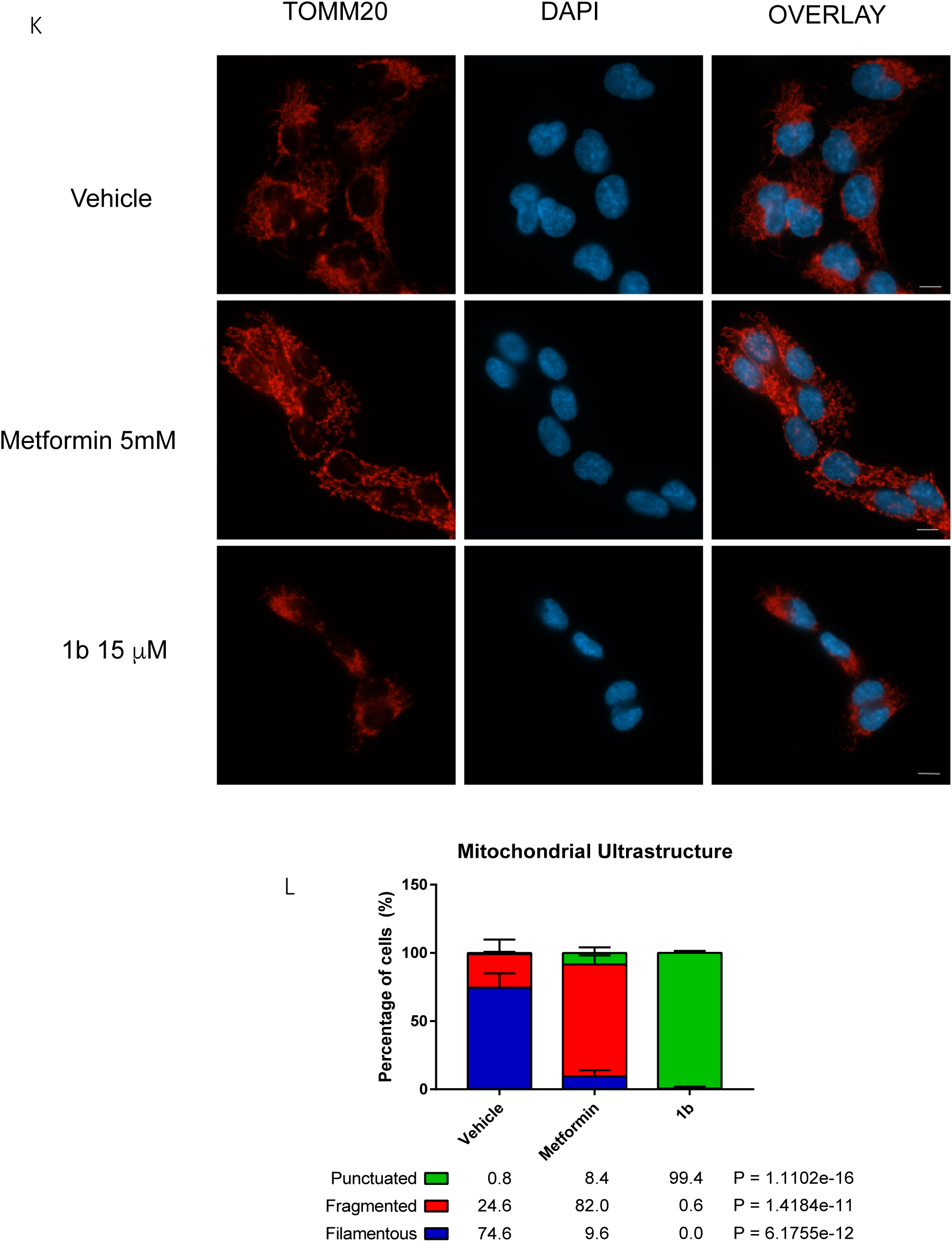
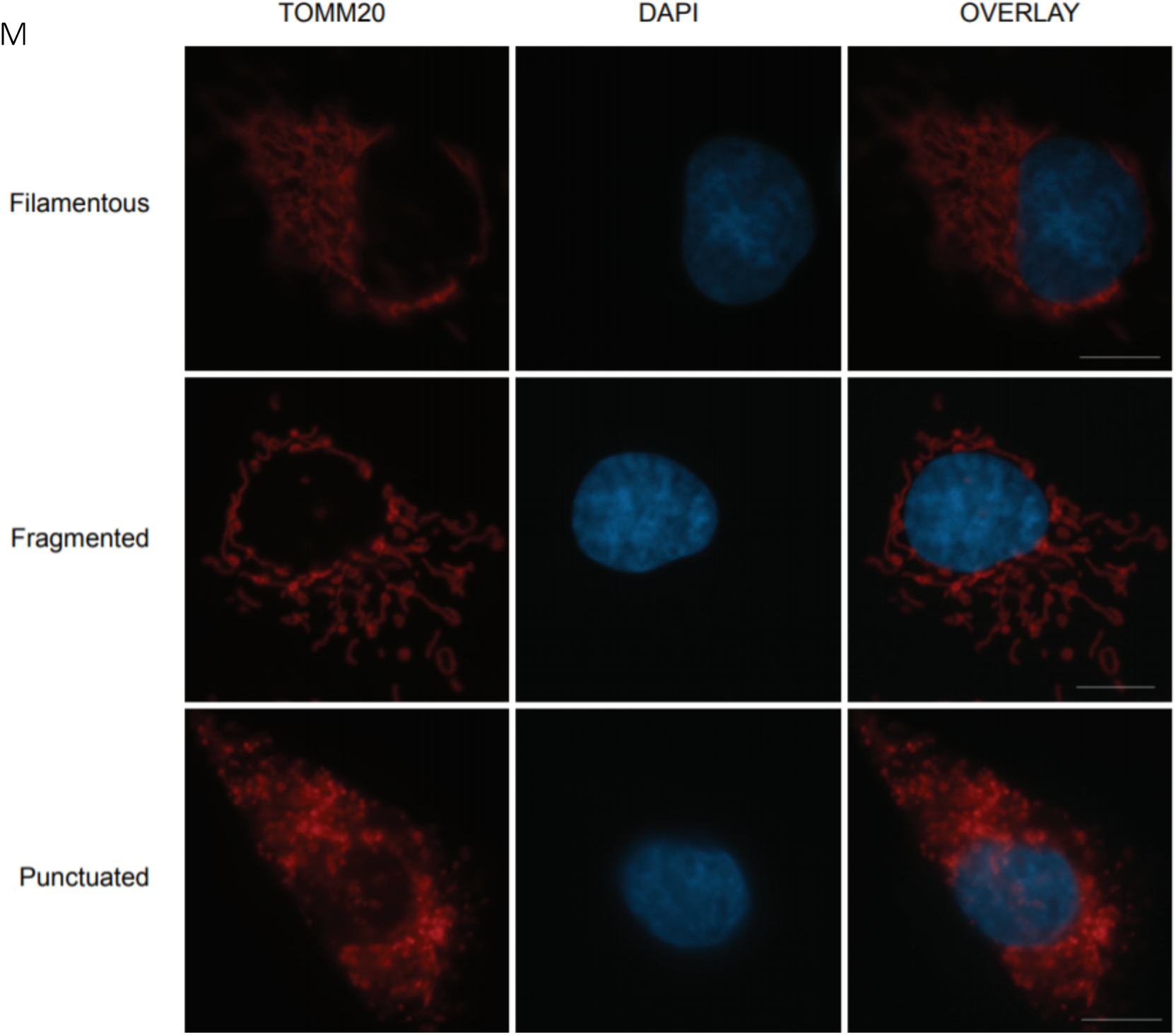
**A) IC_50_ of metformin, phenformin and compound 1b performed *in vitro* over 3 days on KP4 cells. B) IC_50_ of metformin, phenformin and compound 1b performed *in vitro* over 3 days on Panc1 cells**. **C) Effect of 24h treatment with metformin (1 mM) or compound 1b (5 µM) on the formation of tumorspheres in AH375 cells grown in suspension (mouse pancreatic ductal adenocarcinoma).** **** p≤0.0001 (ANOVA). **D) Effect of treatment for 6h of PSN1 cells with metformin (10 mM), phenformin (100 µM) or compound 1b (25 µM) on NAD^+^/NADH ratio**. ** p≤0.01, **** p≤0.0001 (ANOVA). **E) Effect of treatment for 18h of KP4 cells with metformin (5 mM), phenformin (50 µM) or compound 1b (15 µM) on NAD+/NADH ratio**. ** p≤0.01, *** p≤0.001 (ANOVA). **F) Effect of treatment for 24h of KP4 cells with metformin (5 mM), phenformin (50 µM) or compound 1b (15 µM) on phosphorylation level of AMPK T172 and ACC S79. G) Effect of treatment for 24h of KP4 cells with metformin (5 mM), phenformin (50 µM) or compound 1b (15 µM) on cell viability. H) Effect of treatment for 24h of HPNE hTERT cells with metformin (5 mM), phenformin (50 µM) or compound 1b (15 µM) on cell viability.** **** p≤0.0001, NS: not significative (ANOVA). **I) Effect of treatment of KP4 cells with metformin (10 mM) or compound 1b (15 µM) on oxygen consumption rate (OCR) measured by Seahorse analysis.** (**J**) **Effect of treatment of KP4 cells with metformin (10 mM) or compound 1b (15 µM) on** ECAR (extracellular acidification rate) measured by Seahorse analysis. **K) Mitochondrial morphology in KP-4 cells 24h following the indicated treatments as visualised by anti-TOMM20 immunofluorescence**. Scale bar = 10 **µ**M. **L) Quantification of the percentage of cells exhibiting filamentous, fragmented or punctuated mitochondria following indicated treatments**. The data represents the mean of 2 biological replicates and for each replicate three counts of 50 cells were done. n = 300 cells per treatment. Data was analyzed with one way Anova followed by Tukey HSD test. **M) Mitochondrial morphology as visualised by immunofluorescence with anti-TOMM20 antibody.** The filamentous structure is from KP-4 cells treated with vehicle, the fragmented structure is from metformin KP-4 cells treated with metformin, and the punctuated structure is from KP-4 cells treated with **1b**. Scale bar = 10 µm.

We have recently shown that a short treatment of 24h with metformin of the mouse PDAC cell line AH375 grown adherent in 2D decreases the ability of this cell line to subsequently grow as spheres in suspension^23^. Here again compound **1b** show a better biological activity than metformin (Figure 3C) as it decreases by 75% the number of AH375 spheres when treated for 24h with 5 µM of it, while metformin at 1 mM results in a less marked decrease of 50%. In agreement with compound **1b** higher ability to inhibit cancer cells, we observed that compound **1b** induces a significant increase (~20 %) of the fraction of dead KP4 cells (Figure 3G) treated at 15 µM for 24h (measured by trypan blue exclusion) while metformin (5 mM) and phenformin (50 µM) had no significant effects. In contrast with its effect in PDAC cells (KP4), compound **1b** did not increase the fraction of dead cells in normal immortalized non-tumorigenic pancreatic human cells HPNE hTERT in the same conditions (Figure 3H), suggesting selectivity towards cancer cells.

To better characterize the molecular mechanism of action of compound **1b** we measured the NAD^+^/NADH ratio as it was shown that metformin decreases this ratio through inhibition of mitochondrial respiration^24^. In agreement, both in pancreatic cancer PSN1 cells (Figure 3D) and KP4 cells (Figure 3E) compound **1b** at 15 µM decreased NAD^+^/NADH ratio by 25 % and 40 % respectively, as did phenformin (50 µM) and metformin (5 mM). These results suggest that, as metformin and phenformin do, compound **1b** is able to inhibit mitochondrial respiration, but at a much lower concentration. However, as it is broadly reported for metformin and phenformin ^25^, compound **1b** does not activate phosphorylation of AMPK on T172 and that of the AMPK-target ACC (acetyl-CoA carboxilase (Figure 3F).

Next, we used the Seahorse analyser to quantify the ability of compound **1b** to affect oxygen consumption rate (OCR) in KP4 cells. We were able to determine that compound **1b** inhibited OCR at 15 µM, similarly to metformin at 5 mM (Figure 3I). Inhibition of OCR by metformin was simultaneously associated with an increase of the ECAR (roughly: glycolysis), which was also observed with compound **1b** (Figure 3J). These data suggest that the anticancer activity of compound **1b** is due at least in part to the inhibition of mitochondrial respiration, similar to metformin, but at much lower concentration. To further confirm this idea we visualized mitochondria in cells treated with biguanides. Both compound **1b** and metformin led to a lost of the filamentous mitochondrial network but compound **1b** had a bigger impact in mitochondrial morphology leading to a punctuated pattern typical of very dysfunctional mitochondria (Figure 3K-M). Together these data show that compound **1b** functions as metformin or phenformin but with a much higher activity, presumably due to its intrinsic chemical characteristics and membrane activity.

### Antiproliferative activity *in vivo*

To determine whether coumpound **1b** is also superior to phenformin *in vivo*, mice were engrafted with subcutaneous KP4 PDAC cells tumors. After randomization, mice received either phenformin (n = 6 mice, 2 tumors per mouse), compound **1b** (n = 17 mice, 2 tumors per mouse) both at (50 mg/kg/d IP, 5 d/week) or vehicle (n = 10 mice, 2 tumors per mouse). Compound **1b** significantly reduced tumor volume (Figure 4) and mice survival time, while phenformin administred at the same dose had only modest or non-significant effects. Together our data show that in vivo, compound **1b** is well tolerated and more effective than the most active biguanide used curently in human, phenformin.

**Figure 4.**
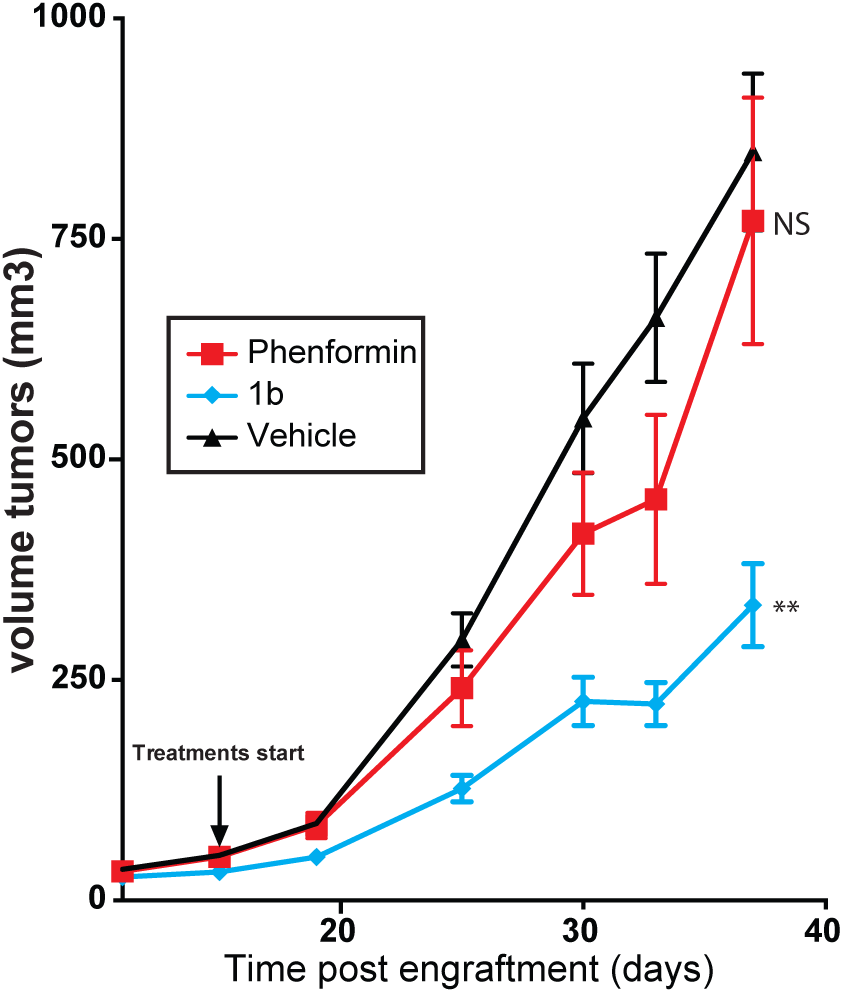
Progression of the volume of KP4 cells sub-cutaneous xenografts performed in nude mice over 37 days. Treatments with phenformin or compound **1b** (both at 50 mg/kg/d, 5 days a week) or vehicle were started 11 days post engrafment. ** p≤0.01, NS: not significative (ANOVA).

## Conclusion

In conclusion, by screening a small library of phenylethynylbenzyl-modified metformin analogs we identified a compound able to penetrate mitochondrial membranes, 1000-fold more active than metformin. Compound **1b** selectively induces the death of PDAC cells (KP4), at least in part by the inhibition of mitochondrial respiration, in a similar way to metformin but at much lower concentration. Compound **1b** is more active than metformin and phenformin to inhibit the proliferation of PDAC cells *in vivo*, as it significantly inhibitis the growth of xenografts and increases mice survival time. We demonstrate herein that the anticancerous properties of biguanides can be improved by chemical modifications. Thenew PEB-Biguanidium compound constitutes a first-class lead compound that can be used to target mitochondria in pancreatic cancer and potentially other cancers.

## Author Contributions

The manuscript was written through contributions of all authors. All authors have given approval to the final version of the manuscript. A.H. prepared figures 1 A-F, figures 2 A-C, M.P. prepared figures 3 A-F, I-J and figure 4, M.-C.R. prepared figures 3 G-M and A.D. prepared figure 2D. All authors declare no conflict of interest.

## Acknowledgments

We gratefully acknowledge the Canadian Institutes of Health Research (CIHR), Canadian Cancer Society (CCS), the Cancer Research Society, Natural Sciences and Engineering Research Council of Canada (NSERC), Fonds Québécois de la Recherche sur la Nature et les Technologies (FRQ-NT), and Fonds Marguerite Ruel pour la Recherché sur le Cancer – Université de Montréal for financial support. G.F. is supported by the CIBC chair for breast cancer research at the CRCHUM.

