## Supplementary material for "Phenylethynylbenzyl-Modified Biguanides Inhibit Pancreatic Cancer Tumor Growth": suplemental file

### 1. Synthesis and Characterization

#### General

All chemicals were purchased from Aldrich Chemicals in their highest purity and used without further purification. Deuterated dimethylsulfoxide (DMSO-$d^6$) and deuterated chloroform ($CDCl_3$) was purchased from CDN Isotopes. NMR spectra were recorded on a Bruker advance 400. Coupling constants are given in hertz (Hz) and chemical shifts are given in parts per million (ppm, δ) measured relative to the residual solvent (the multiplicity of the signals are given as s: singlet, d: doublet, t: triplet, and m: multiplet). High-resolution mass spectra (HRMS) were recorded on a TSQ Quantum Ultra (Thermo Scientific) triple quadrupole with accurate mass option instrument. (Université de Montréal Mass Spectrometry Facility). MS experiences were performed using an UltrafleXtreme MALDI TOF/TOF mass spectrometer equipped with a SmartBeam II Nd:Yag/355 nm laser operating at 1 kHz and providing a laser focus down to 20 µm in diameter (Bruker Daltonics, Billerica, MA). The data acquisition for MS was performed in positive ion mode using the linear geometry with flexControl 3.4 (Bruker Daltonics, Billerica, MA). Acceleration voltage was set to +25kV and all other instrumental parameters (delayed extraction parameters, source voltages, detector gain, laser energy, etc.) were optimized for maximum S/N for the drug compounds. L-α-Phosphatidylcholine was purchased from Avanti Polar Lipids. Transport and depolarization studies as well as absorbance measurements were performed on a Varian Cary Eclipse fluorescence spectrophotometer. The hemolysis assay was performed on a Fluostar Optima plate reader.

#### Synthetic procedures

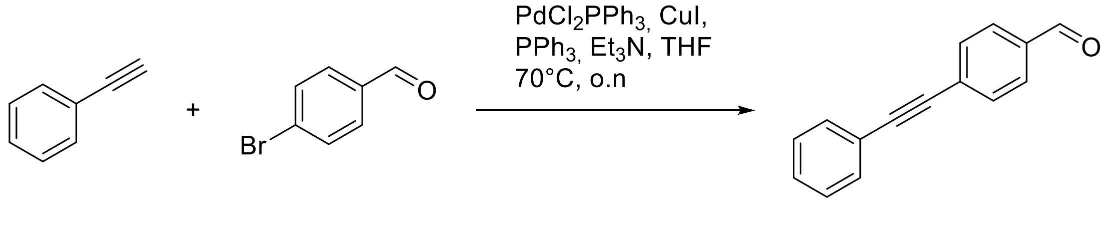

***4-(phenylethynyl)benzaldehyde.*** 4-Bromobenzaldehyde (4.00 g, 21.6 mmol), $PdCl_2(PPh_3)_2$ (0.091g, 0.13 mmol), CuI (0.082g, 0.43 mmol) and triphenylphosphine (0.113g, 0.43 mmol) were dissolved in a mixture of THF (100 ml) and triethylamine (22.4 ml, 173 mmol). Phenylacetylene (2.85 ml, 25.9 mmol) was added slowly and the mixture was heated to 70°C overnight. The mixture was cooled to room temperature, filtered and washed with THF and concentrated in vacuo. The residue was then purified by flash chromatography (EtOAc/hexane gradient) to afford 4-(phenylethynyl)benzaldehyde (4.42 g, 21.6 mmol) as a crystalline solid (quantitative yield %)

$^1$H NMR (400 MHz, $CDCl_3$, 25 °C, TMS): δ = 7.40 (m, 3 H), 7.58 (m, 2H), 7.70 (d, *J* = 8.0 Hz, 2H), 7.90 (d, *J* = 4.0 Hz, 2H), 10.05 (s, 1H)

$^{13}$C NMR (70 MHz, $CDCl_3$, 25 °C, TMS,): δ = 89.1, 93.4, 122.1, 128.6, 129.3, 129.9, 130.1, 132.1, 132.5, 135.9, 192.9

HRMS (120.0V, ES+): *m*/*z* (%) = 207.08130 ($[C_{15}H_{11}N_5]$+H)$^+$, 208.0843 (M + H)$^+$.

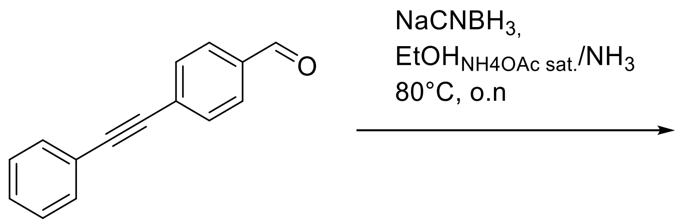

***4-(phenylethynyl)benzylamine.*** 4-(phenylethynyl)benzaldehyde (4.42g, 21.4 mmol) and sodium cyanoborohydride (4.04, 64.3 mmol) were dissolved in a solvent mixture of EtOH saturated with $NH_4OAc$ and $NH_4OH$ 5:2 (30 mM), and reaction was heated at 80°C overnight. EtOH was evaporated under reduced pressure, resulting mix was extracted with DCM and washed with $NaHCO_3$. The mixture was concentrated in vacuo and purified by flash chromatography (DCM/MeOH gradient) to afford 4-(phenylethynyl)benzylamine (1.02 g, 4.9 mmol) as a white solid. (20% yield)

$^{1}$H NMR (400 MHz, DMSO-$d_6$, 25 °C, TMS): δ = 3.77 (s, 2H), 7.42 (m, 6H), 7.52 (d, $J$ = 8.0 Hz, 2H), 7.55 (m, 3H)

$^{13}$C NMR (70 MHz, DMSO-$d_6$, 25 °C, TMS): δ = 42.4, 89.8, 90.4, 122.5, 122.8, 129.3, 129.4, 129.7, 131.9, 132.0, 135.1

HRMS (120.0V, ES+): $m/z$ (%) = 208.11260 ($[C_{15}H_{14}N_1]+H)^+$, 209.1163 $(M + H)^+$.

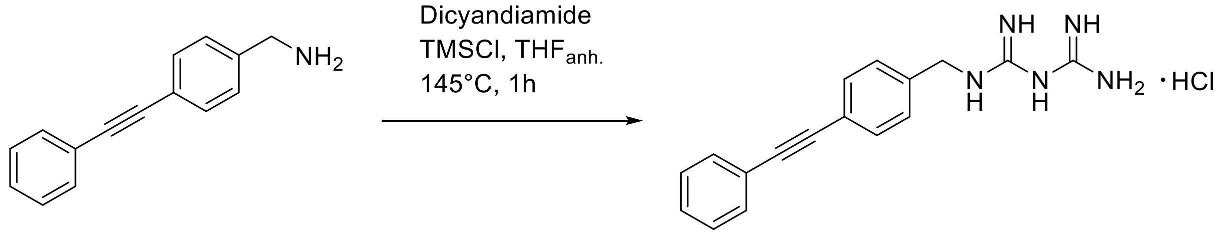

***4-(phenylethynyl)benzylbiguanide chloride salt (1).*** 4-(phenylethynyl)benzylamine (1.00 g, 4.82 mmol), dicyandiamide (0.811 g, 9.64 mmol) and trimethylsilylchloride (2.45 ml, 19.3 mmol) were dissolved in anhydrous THF (23.8ml, 202 mM) in a sealed tube. The mixture was heated to 145°C for 1h. The mixture was filtered and washed with THF, and residue was purified by TLC prep (DCM:MeOH 9:1) to afford 4-(phenylethyl)benzylbiguanide as its chloride salt (0.460g, 1.40 mmol, 30% yield)

$^{1}$H NMR (400 MHz, DMSO-$d_6$, 25 °C, TMS): δ = 4.41 (d, 2H), 7.00 (s, 6H), 7.37 (d, $J$ = 4.0 Hz 3H), 7.44 (m, 4H), 7.55 (m, 5H)

$^{13}$C NMR (70 MHz, DMSO-$d_6$, 25 °C, TMS): δ = 44.5, 89.6, 121.4, 122.7, 128.0, 129.2, 131.8, 158.9, 160.8

HRMS (120.0V, ES+): $m/z$ (%) = 292.1567($[C_{17}H_{22}N_5]+H)^+$, 293.1590 $(M + H)^+$.
Purity was assessed by HPLC, > 99%

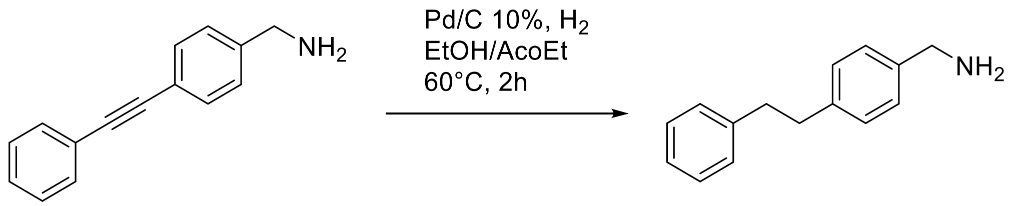

***4-(phenylethyl)benzylamine.*** 4-(phenylethynyl)benzylamine (500 mg, 2.41 mmol) and palladium on carbon 10 wt.% (0.51g, 0.48 mmol) were mixed in 200 ml of a EtOH:AcOEt 1:1 mixture under nitrogen. Reaction was put in a $H_2$ atmosphere and heated to 60 °C for 2h. After purging with nitrogen, reaction was filtered on a celite pad and evaporated under reduced pressure to afford 4-(phenylethyl)benzylamine (500 mg, 2.41 mmol) as a white solid (quantitative yield)

$^{1}$H NMR (400 MHz, DMSO-$d_6$, 25 °C, TMS): δ = 2.87 (s, 4H), 3.70 (s, 2H), 7.18 (m, 3H), 7.27 (m, 8H)

$^{13}$C NMR (70 MHz, DMSO-$d_6$, 25 °C, TMS): δ = 37.2, 37.6, 45.5, 126.3, 127.6, 128.6, 128.7, 128.9, 140.0, 141.2, 142.0

LRMS (120.0V, ES+): *m/z* (%) = 212.14 ($[C_{15}H_{19}N_1]$+H)$^+$

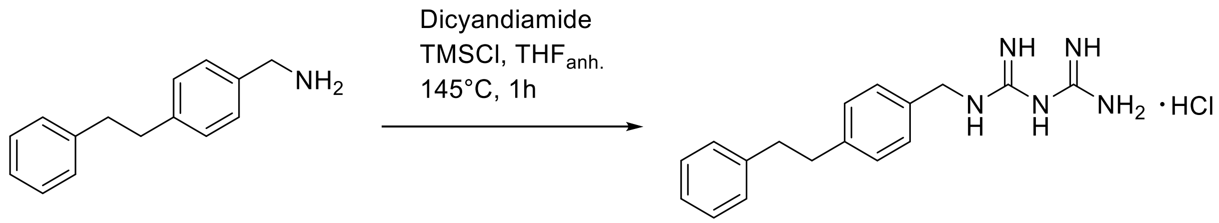

***4-(phenylethyl)benzylbiguanide chloride salt (2).*** 4-(phenylethyl)benzylamine (0.273 g, 1.29 mmol), dicyandiamide (0.217 g, 2.58 mmol) and trimethylsilylchloride (0.655 ml, 5.16 mmol) were dissolved in anhydrous THF (6.38 ml, 202 mM) in a sealed tube. The mixture was heated to 145°C for 1h. The mixture was filtered and washed with THF, and residue was purified by TLC prep (DCM:MeOH 9:1) to afford 4-(phenylethyl)benzylbiguanide as its chloride salt (72 mg, 0.24 mmol, 20% yield)

1H NMR (400 MHz, DMSO-$d_6$, 25 °C, TMS): $\delta$ = 2.87 (s, 4H), 4.30 (d, *J* = 4.0 Hz, 2H), 6.92 (s, 5H), 7.24 (m, 10H), 7.64 (s, 1H)

$^{13}$C NMR (70 MHz, DMSO-$d_6$, 25 °C, TMS): $\delta$ = 37.2, 37.5, 42.5, 44.5, 126.3, 127.7, 128.7, 128.8, 129.0, 129.4, 132.0, 140.7, 141.8, 141.9, 142.3, 159.0, 160.3

HRMS (120.0V, ES+): *m/z* (%) = 296.1876 ($[C_{17}H_{22}N_5]$+H)$^+$
Purity was assessed by HPLC, 95%

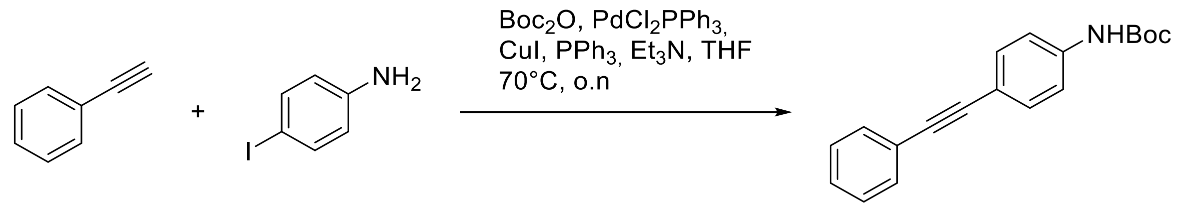

***tert-butyl (4-(phenylethynyl)phenyl)carbamate.*** Iodoaniline (3.0 g, 14.0 mmol) and di-*tert*-butyl dicarbonate (3.0 g, 14.0 mmol) were mixed in THF at room temperature overnight. $PdCl_2(PPh_3)_2$ (0.058g, 0.082 mmol), CuI (0.047g, 0.28 mmol) and triphenylphosphine (0.047g, 0.28 mmol) and triethylamine (13.5 ml, 112.0 mmol) were added to the mix, and phenylacetylene (1.5 ml, 14.0 mmol) was added slowly. Reaction was heated to 60°C ovenight. The mixture was cooled to room temperature, filtered and concentrated in vacuo. The residue was then purified by flash chromatography (EtOAc/hexane gradient) to afford *tert*-butyl (4-(phenylethynyl)phenyl)carbamate (1.39 g, 4.7 mmol, 78% yield)

$^1$H NMR (400 MHz, $CDCl_3$, 25 °C, TMS):δ = 1.55 (s, 9 H), 6.57 (s, 1H), 7.37 (m, 5H), 7.49 (d, *J* = 8.0 Hz, 2H), 7.54 (m, 2H)

$^{13}$C NMR (70 MHz, $CDCl_3$, 25 °C, TMS): δ = 27.4, 28.3, 89.3, 117.5, 118.03, 123.43, 128.0, 128.3, 131.5, 132.45, 138.4, 146.75

LRMS (120.0V, ES+): *m/z* (%) = 294.38 ($[C_{19}H_{19}NO_2]$+H)$^+$

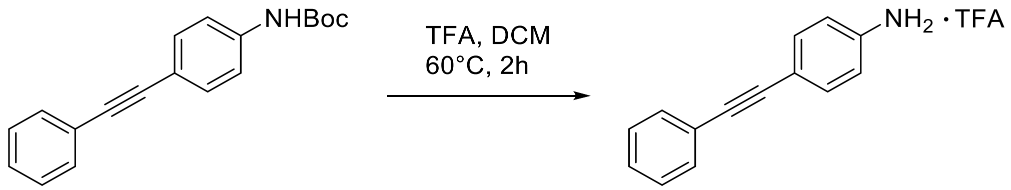

***4-(phenylethynyl)aniline trifluoroacetic salt.*** *tert*-butyl (4-(phenylethynyl)phenyl)carbamate (1.39g, 4.7 mmol) and trifluoroacetic acid (5 eq.) were stirred in DCM at 60 °C for 2h. Solvent was evaporated under reduced pressure and reaction was purified on a silica column (DCM:MeOH gradient) to afford 4-(phenylethynyl)aniline trifluoroacetic salt (1.44 g, 4.7 mmol, quantitative yield)

$^{1}$H NMR (400 MHz, DMSO-d$_6$, 25 °C, TMS): δ = 4.15 (s, 2 H), 6.57 (d, $J$ = 8.0 Hz, 2H), 7.27 (m, 5H), 7.77 (d, $J$ = 12.0 Hz, 2H)

$^{13}$C NMR (70 MHz, DMSO-d$_6$, 25 °C, TMS): δ = 91.6, 114.1, 123.8, 128.0, 129.1, 131.2, 133.2, 150.0

LRMS (120.0V, ES+): $m/z$ (%) = 212.11 ($[C_{14}H_{11}N_1]$+H)$^+$

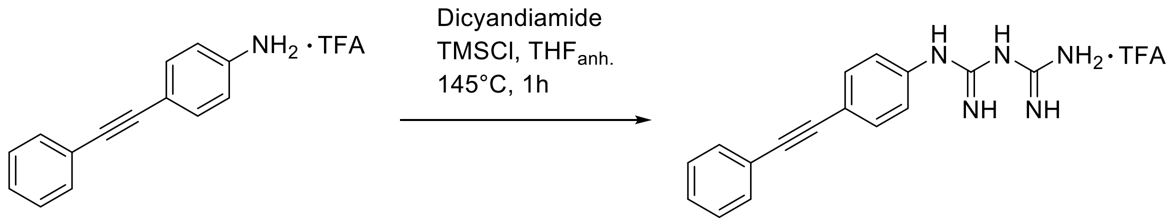

***(4-(phenylethynylphenyl)biguanide trifluoroacetic salt (3).*** 4-(phenylethynyl)aniline trifluoroacetic salt (0.350 g, 1.19 mmol), dicyandiamide (0.20 g, 2.37 mmol) and trimethylsilylchloride (0.602 ml, 4.74 mmol) were dissolved in anhydrous THF (5.86 ml, 202 mM) in a sealed tube. The mixture was heated to 145°C for 1h. The mixture was filtered and washed with THF, and residue was triturated with EtOH to afford (4-(phenylethynylphenyl)biguanide as its trifluoroacetic acid salt (0.18 g, 0.31 mmol, 26% yield)

$^{1}$H NMR (400 MHz, DMSO-d$_6$, 25 °C, TMS): $\delta$ = 7.08 (s, 3 H), 7.27 (m, 6H), 7.45 (s, 3H), 7.52 (d, $J$ = 8.0 Hz, 2H), 7.99 (d, $J$ = 12.0 Hz, 2H)

$^{13}$C NMR (70 MHz, DMSO-d$_6$, 25 °C, TMS): $\delta$ = 44.9, 119.5, 126.9, 128.7, 130.0, 131.0, 135.9, 156.0, 163.3

MS (120.0V, ES+): $m/z$ (%) = 296.1508 ([$C_{16}H_{15}N_5$]+H+$NH_4$)$^+$
Purity was assessed by HPLC, 95%

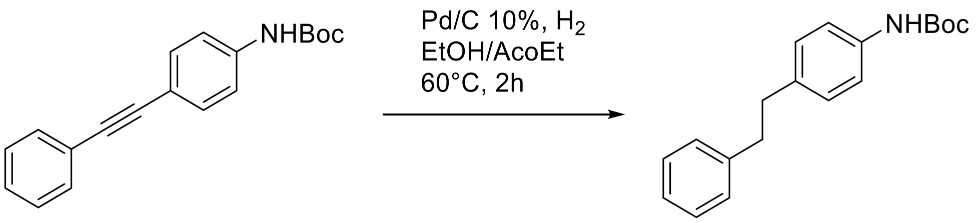

***tert-butyl (4-phenethylphenyl)carbamate.*** *tert*-butyl (4-(phenylethynyl)phenyl)carbamate (0.35g, 1.19 mmol) and palladium on carbon 10 wt.% (0.25 g, 0.239 mmol) were mixed in 200 ml of a EtOH:AcOEt 1:1 mixture under nitrogen. Reaction was put in a $H_2$ atmosphere and heated to 60 °C for 2h. After purging with nitrogen, reaction was filtered on a celite pad and evaporated under reduced pressure to afford *tert*-butyl (4-phenethylphenyl)carbamate (0.35g, 1.9 mmol) as a white solid (quantitative yield)

$^1$H NMR (400 MHz, $CDCl_3$, 25 °C, TMS): δ = 1.53 (s, 9H),2.89 (s, 4H), 6.42 (s, 1H), 7.11 (d, $J$ = 8.0 Hz, 3H), 7.19 (m, 4H) ,7.28 (m, 2H)

$^{13}$C NMR (70 MHz, $CDCl_3$, 25 °C, TMS): δ = 28.5, 37.3, 38.1, 58.5, 80.4, 118.7, 126.0, 128.3, 128.6, 129.0, 136.3, 136.6, 141.9

LRMS (120.0V, ES+): $m/z$ (%) = 298.52 ($[C_{19}H_{23}NO_2]+H)^+$

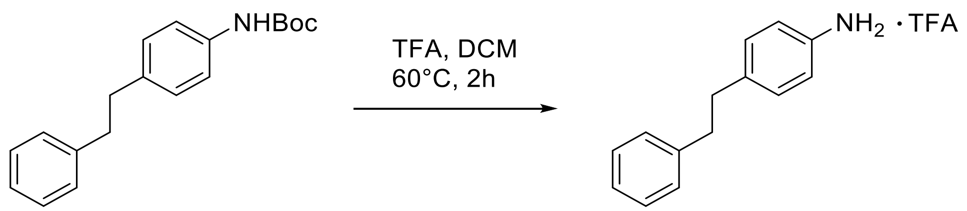

***4-(phenylethylphenyl)aniline trifluoroacetic salt.*** *tert*-butyl (4-(phenylethyl)phenyl)carbamate (0.2 g, 0.63 mmol) and trifluoroacetic acid (5 eq.) were stirred in DCM at 60 °C for 2h. Solvent was evaporated under reduced pressure and reaction was purified on a silica column (DCM:MeOH gradient) to afford 4-(phenylethylphenyl)aniline trifluoroacetic salt (1.44 g, 4.7 mmol, quantitative yield)

$^{1}$H NMR (400 MHz, DMSO-d$_6$, 25 °C, TMS): δ = 2.88 (s, 4 H), 7.07 (s, 1H), 7.18 (m, 6H), 7.27 (m, 5H),

$^{13}$C NMR (70 MHz, DMSO-d$_6$, 25 °C, TMS): δ = 36.6, 37.4, 121.9, 126.3, 128.7, 128.9, 130.0, 133.0, 141.7

LRMS (120.0V, ES+): *m/z* (%) = 198.50 ([$C_{14}H_{15}N_1$]+H)$^+$

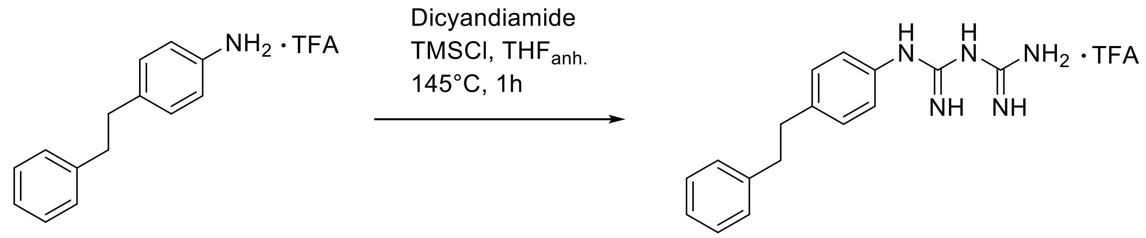

***(4-(phenylethylphenyl)biguanide trifluoroacetic acid salt (4).*** 4-(phenylethylphenyl)aniline trifluoroacetic salt (0.350 g, 1.17 mmol), dicyandiamide (0.20 g, 2.34 mmol) and trimethylsilylchloride (0.594 ml, 4.68 mmol) were dissolved in anhydrous THF (5.86 ml, 202 mM) in a sealed tube. The mixture was heated to 145°C for 1h. The mixture was filtered and washed with THF, and residue was triturated with EtOH to afford (4-(phenylethylphenyl)biguanide as its trifluoroacetic acid salt (0.150 g, 0.41 mmol, 35%)

$^1$H NMR (400 MHz, DMSO-$d_6$, 25 °C, TMS): δ = 2.86 (s, 4 H), 7.23 (m, 10H), 7.45 (s, 6H), 8.70 (s, 1H)

$^{13}$C NMR (70 MHz, DMSO-$d_6$, 25 °C, TMS): δ = 36.9, 37.5, 122.2, 126.3, 128.7, 128.9, 129.2, 136.9, 141.9, 155.9, 161.4

MS (120.0V, ES+): *m/z* (%) = 282.1721 ($[C_{16}H_{19}N_5]+H)^+$
Purity was assessed by HPLC, > 99%

***Anion exchange: general procedure.***

PEB-biguanidium **3** and **4** as their trifluoroacetic salt were dissolved in methanol and deprotonated with $NaHCO_3$ (4 equivalents) for 3 hours at room temperature. The mixture was concentrated in vacuo and the residue was triturated with EtOAc. The precipitate was filtered and washed to afford deprotonated PEB-biguanide **3** and **4** in quantitative yield.

PEB-biguanidium **1** and **2** as their chloride salt and bis(trifluoromethane)sulfonamide lithium salt ($LiNTf_2$, 2.5 equivalents) or lithium trifluoromethanesulfonate (LiOTf, 2.5 eq) were dissolved in methanol and stirred overnight at room temperature. PEB-biguanidium

**3** and **4** were prepared following the same procedure with the addition of 2 equivalents of HCl to the reaction. The mixture was concentrated in vacuo and the residue was triturated in EtOAc. The precipitate was then filtered and washed with EtOAc to afford the PEB-biguanidium **1, 2, 3** or **4** as a bis(trifluoromethane)sulfonamide salt or a trifluoromethanesulfonate salt in quantitative yields.

OTF:

MS (120.0V, ES-): *m/z* (%) = 148.9527 ($[CF_3O_3SH]-H)^-$

$^{19}F$ NMR (282 MHz, DMSO-$d_6$, 25 °C, TMS): δ = 77.75 (s, 3F)

$NTf_2$:

MS (120.0V, ES-): *m/z* (%) = 279.9192 ($[C_2F_6O_4S_2H]-H)^-$

$^{19}F$ NMR (282 MHz, DMSO-$d_6$, 25 °C, TMS): δ = 78.73 (s, 6F)

#### NMR Spectra

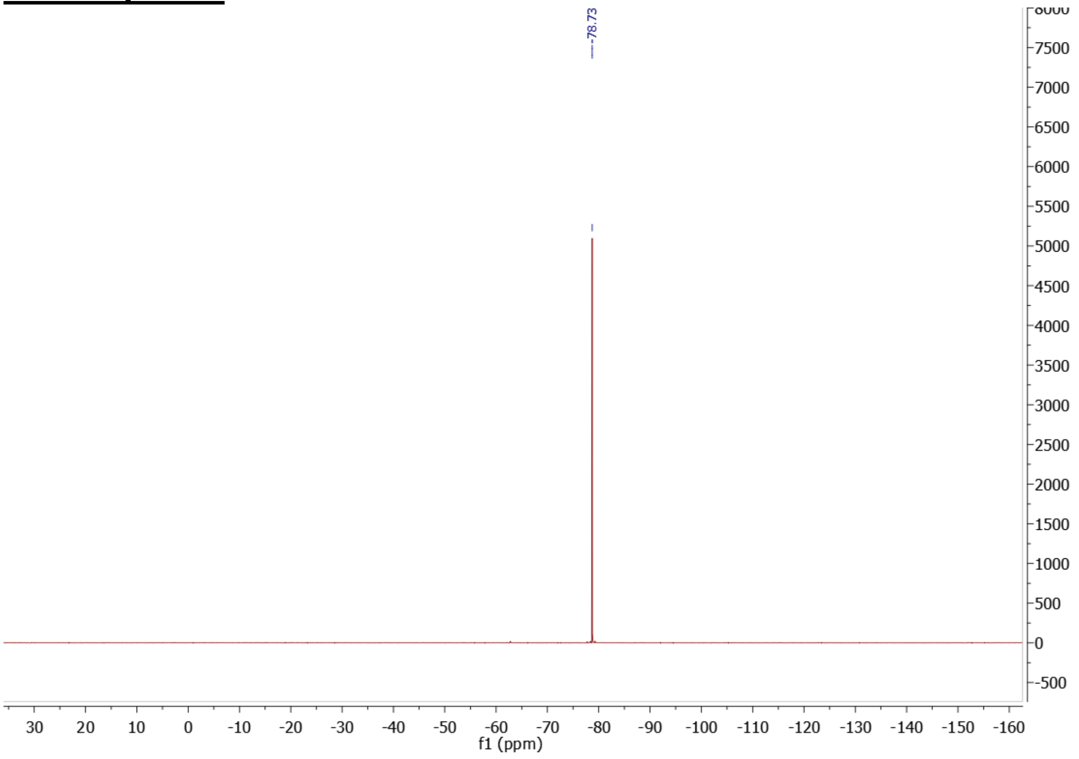

Figure S1: $^{19}F$ NMR (282 MHz) spectrum of $NTf_2^-$ (from 1a) in DMSO-d6 at 298K

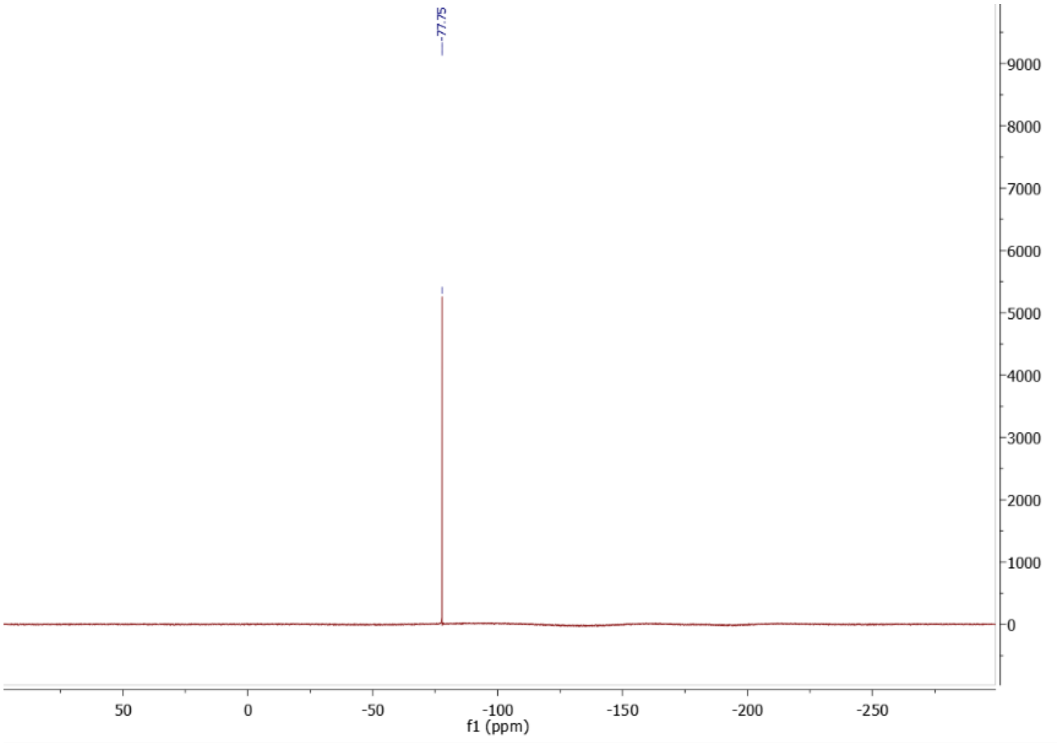

Figure S2: $^{19}$F NMR (282 MHz) spectrum of OTf $^{-}$ (from 1b) in DMSO-d6 at 298K

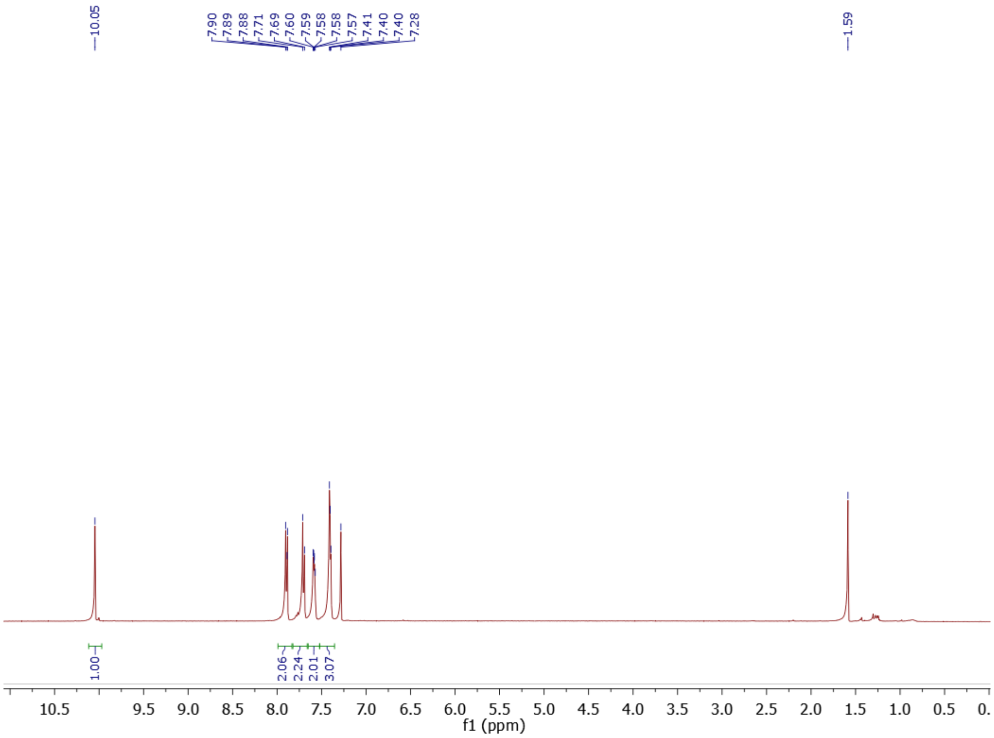

Figure S3: $^{1}$H NMR (400 MHz) spectrum of 4-(phenylethynyl)benzaldehyde in $CDCl_3$ at 298K

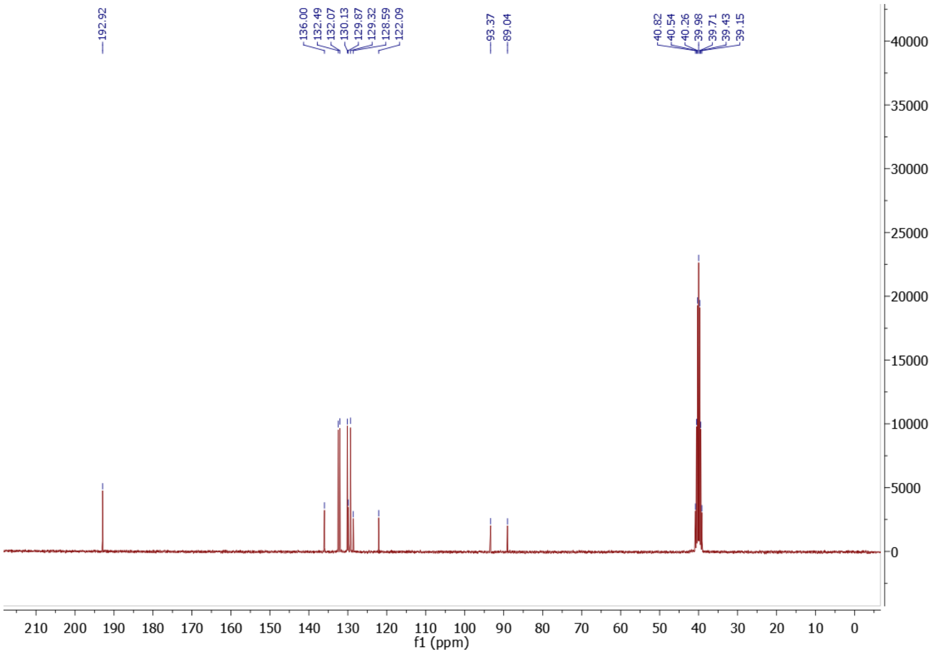

Figure S4: $^{13}$C NMR (70 MHz) spectrum of 4-(phenylethynyl)benzaldehyde in $CDCl_3$ at 298K

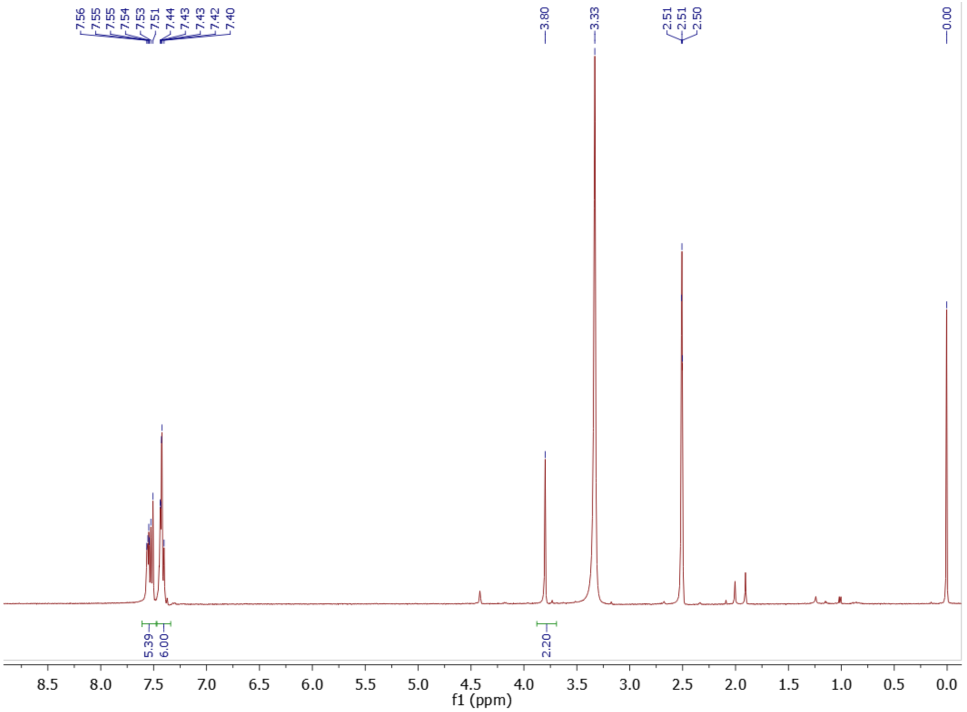

Figure S5: $^{1}$H NMR (400 MHz) spectrum of 4-(phenylethynyl)benzylamine in DMSO-d6 at 298K

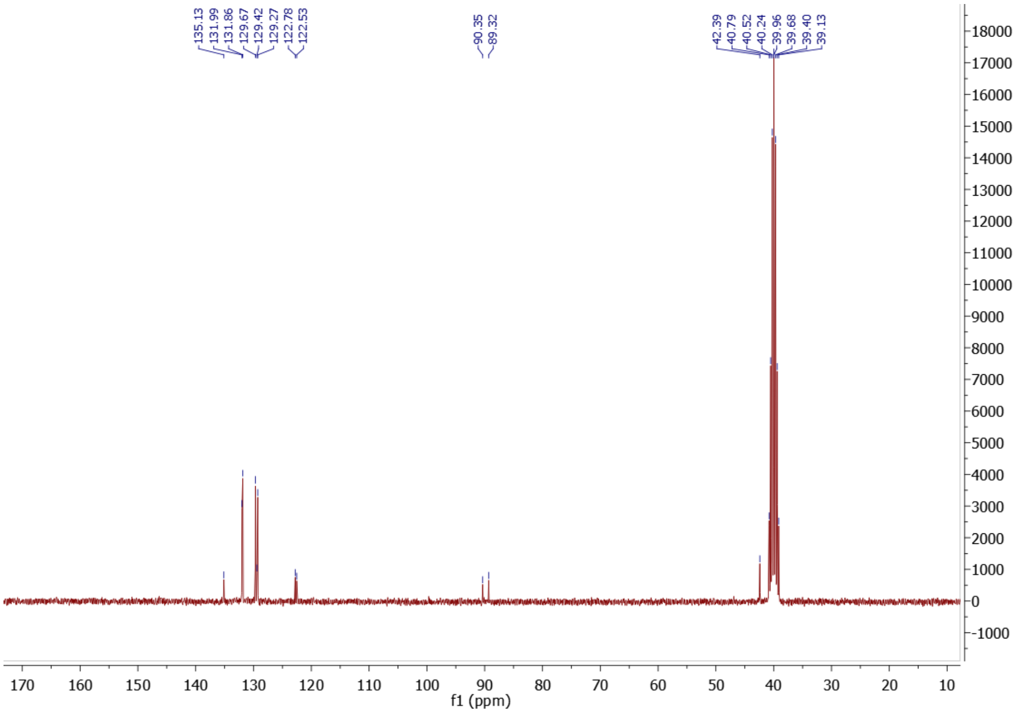

Figure S6: $^{13}$C NMR (70 MHz) spectrum of 4-(phenylethynyl)benzylamine in DMSO-d6 at 298K

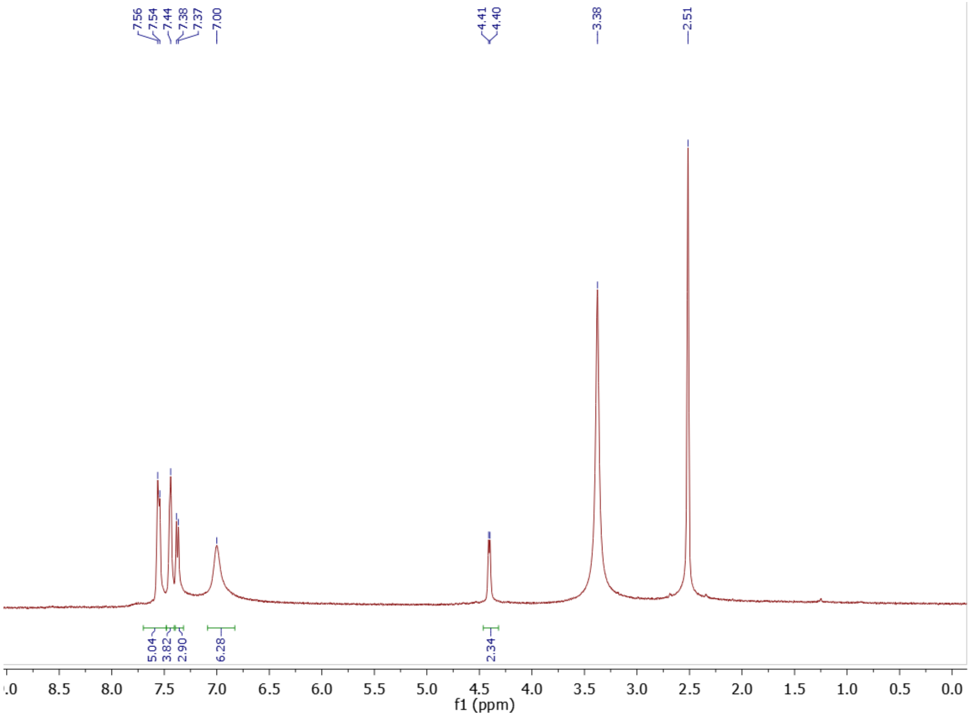

Figure S7: [1]H NMR (400 MHz) spectrum of 4-(phenylethynyl)benzylbiguanide chloride salt **(1)** in DMSO-d6 at 298K

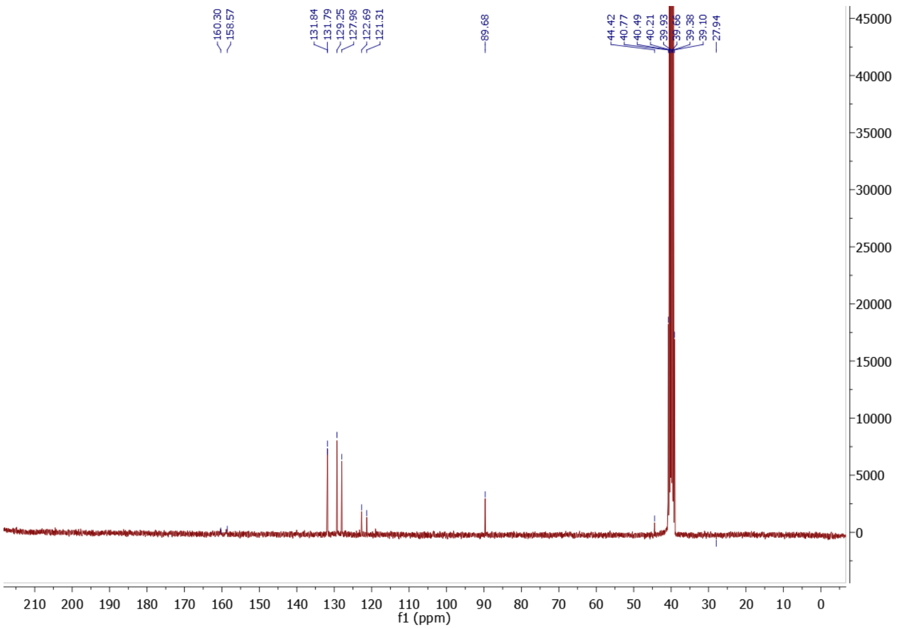

Figure S8: [13]C NMR (70 MHz) spectrum of 4-(phenylethynyl)benzylbiguanide chloride salt **(1)** in DMSO-d6 at 298K

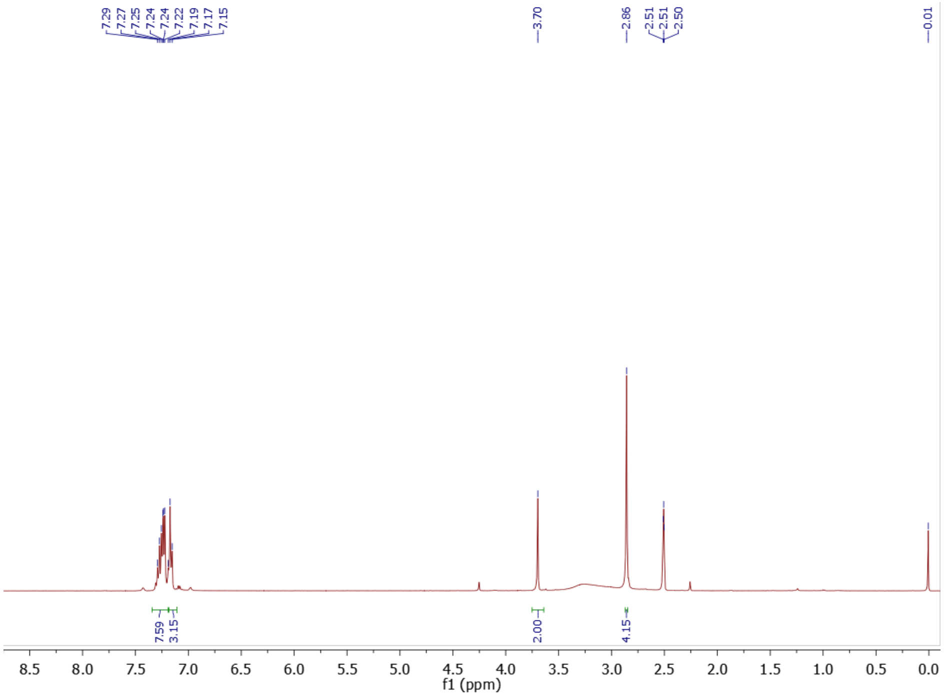

Figure S9: $^{1}$H NMR (400 MHz) spectrum of 4-(phenylethyl)benzylamine in DMSO-d6 at 298K

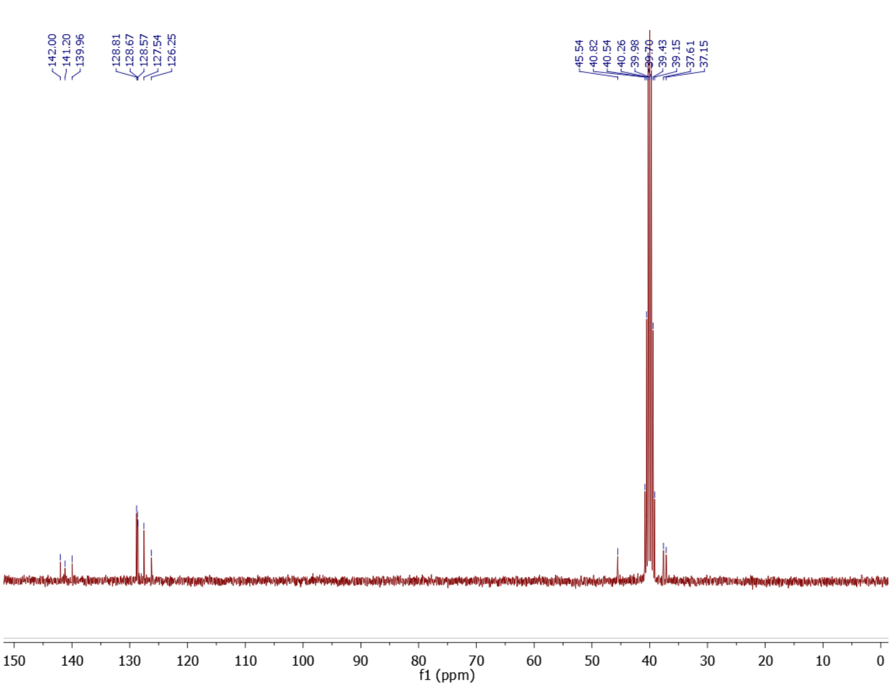

Figure S10: $^{13}$C NMR (70 MHz) spectrum of 4-(phenylethyl)benzylamine in DMSO-d6 at 298K

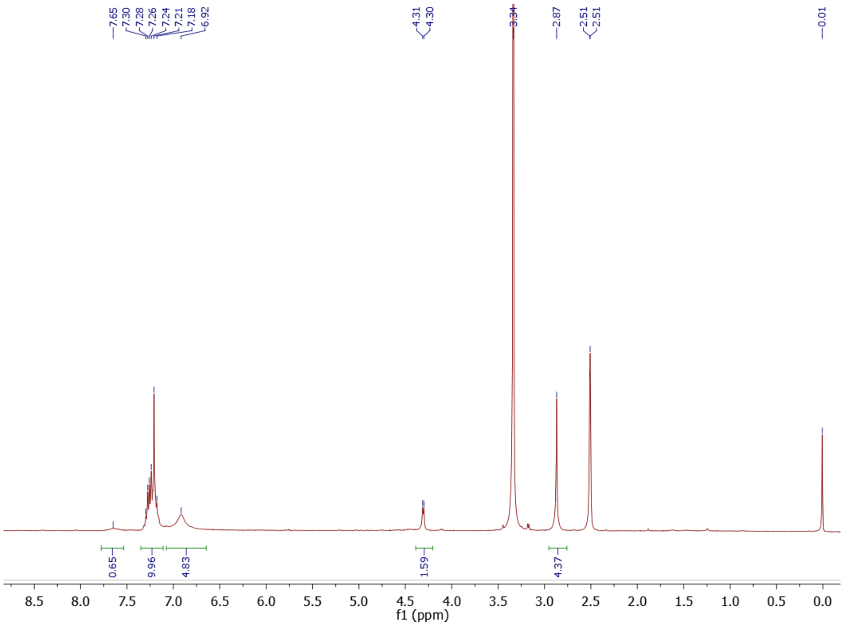

Figure S11: $^{1}$H NMR (400 MHz) spectrum of 4-(phenylethyl)benzylbiguanide chloride salt **(2)** in DMSO-d6 at 298K

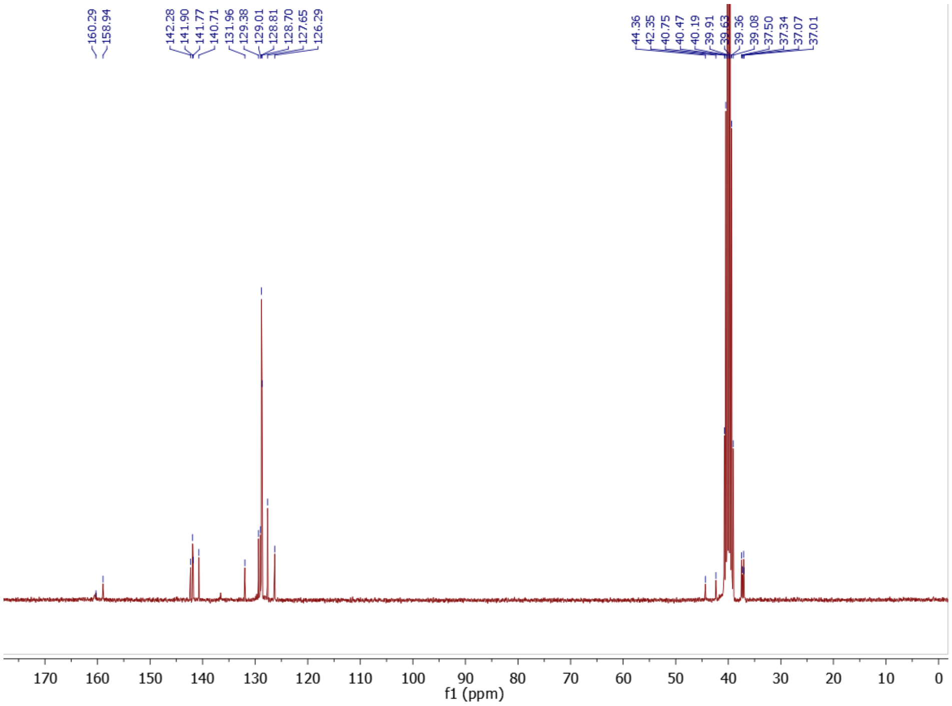

Figure S12: $^{13}$C NMR (70 MHz) spectrum of 4-(phenylethyl)benzylbiguanide chloride salt **(2)** in DMSO-d6 at 298K

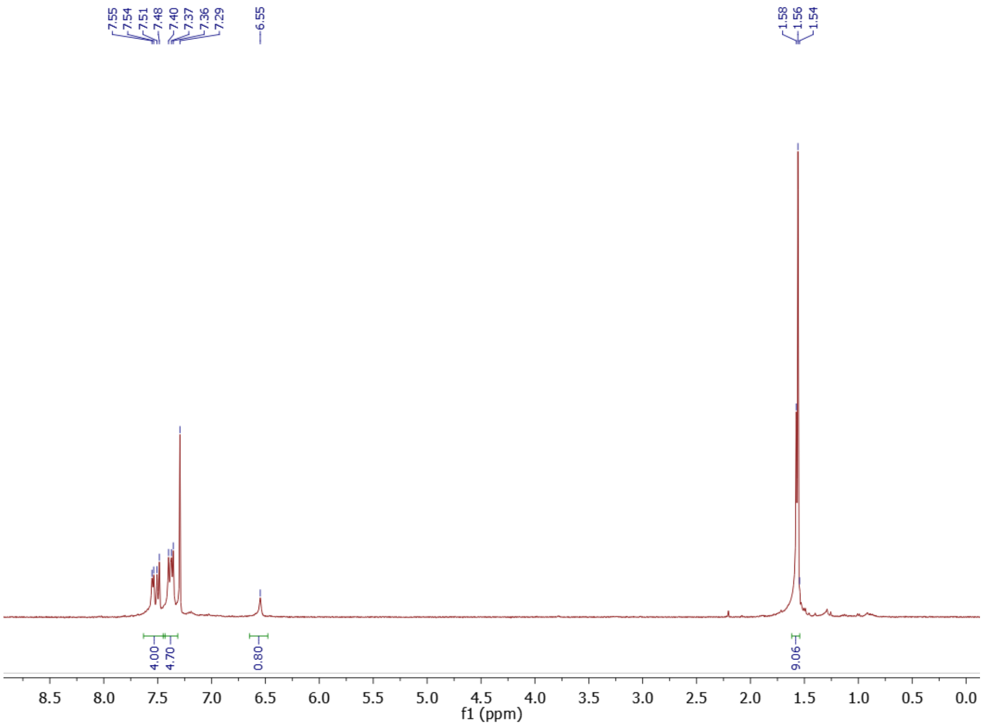

Figure S13: $^{1}$H NMR (400 MHz) spectrum of tert-butyl (4-(phenylethynyl)phenyl)carbamate in $CDCl_3$at 298K

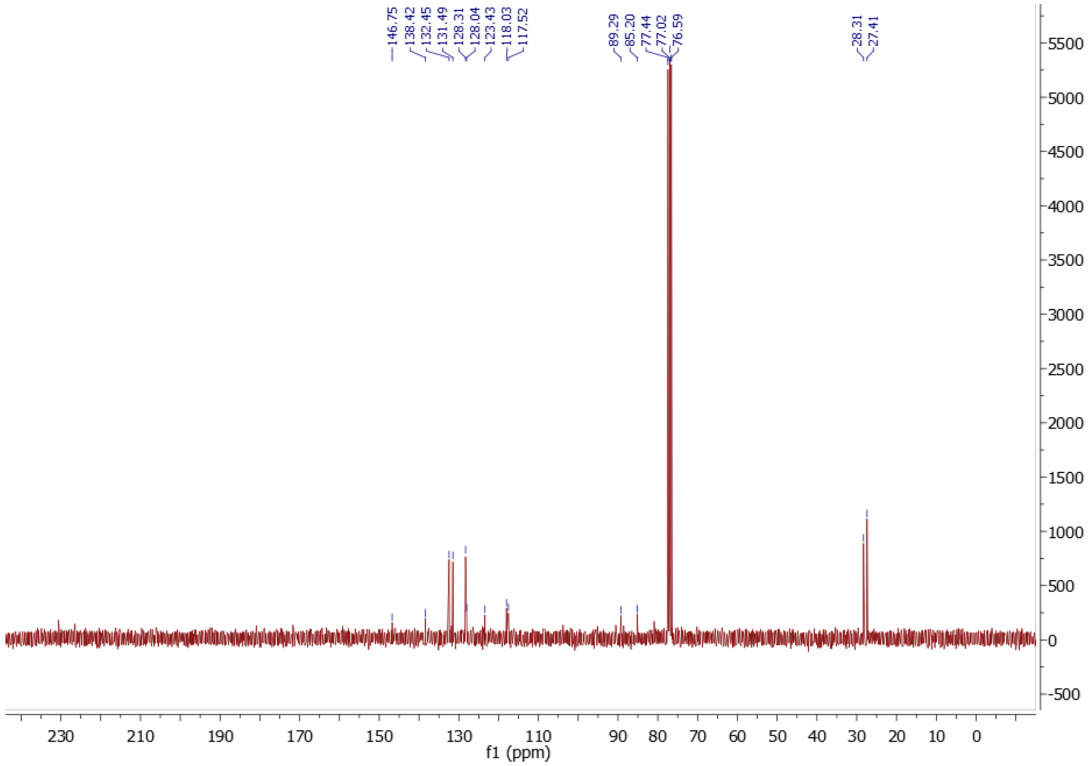

Figure S14: $^{13}$C NMR (70 MHz) spectrum of tert-butyl (4-(phenylethynyl)phenyl)carbamate in $CDCl_3$ at 298K

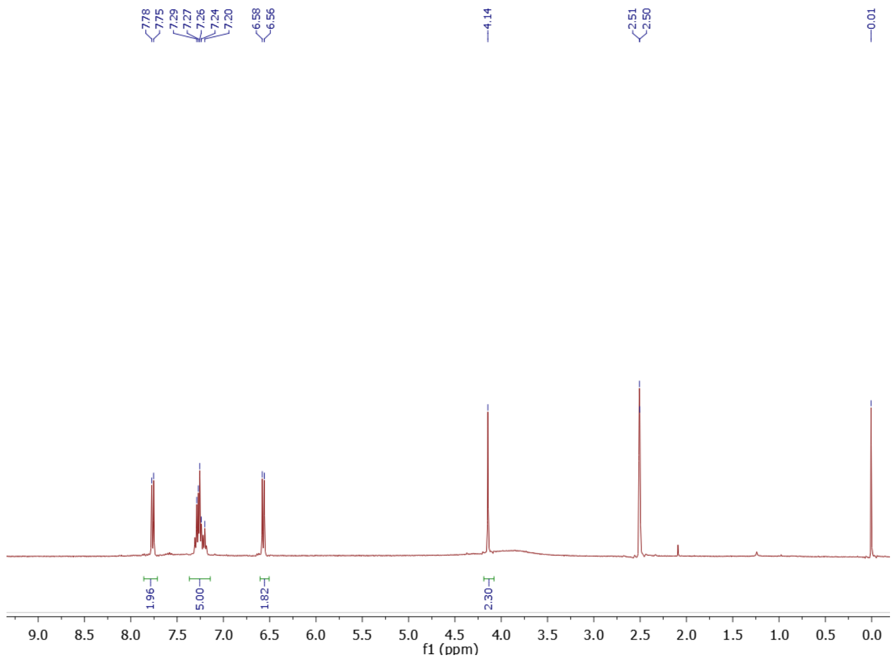

Figure S15: $^{1}$H NMR (400 MHz) spectrum of 4-(phenylethynyl)aniline trifluoroacetic salt in DMSO-d6 at 298K

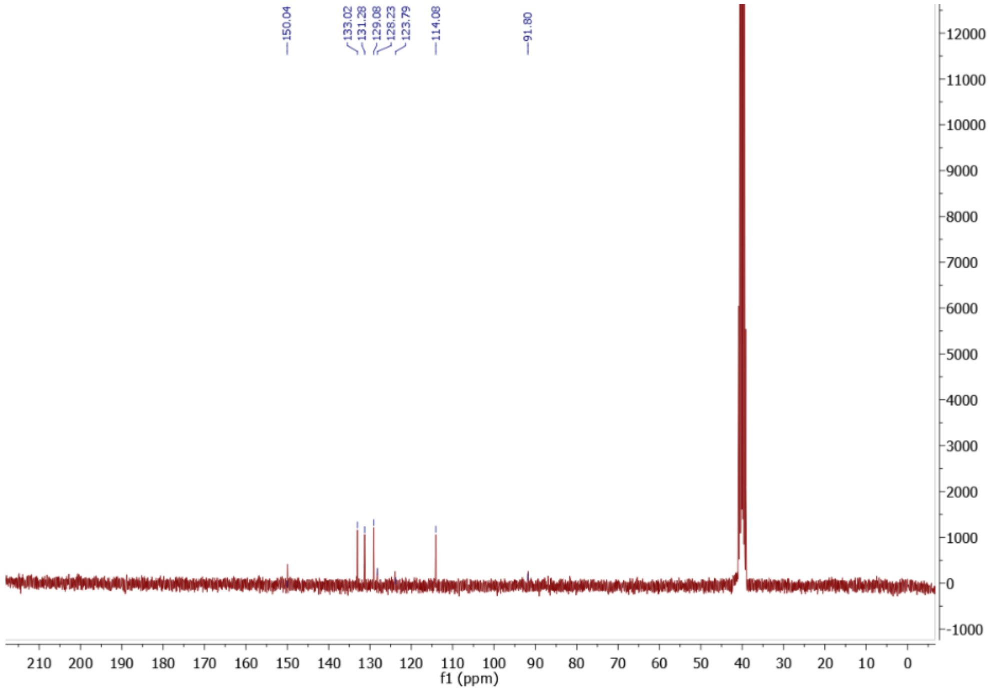

Figure S16: $^{13}$C NMR (70 MHz) spectrum of 4-(phenylethynyl)aniline trifluoroacetic salt in DMSO-d6 at 298K

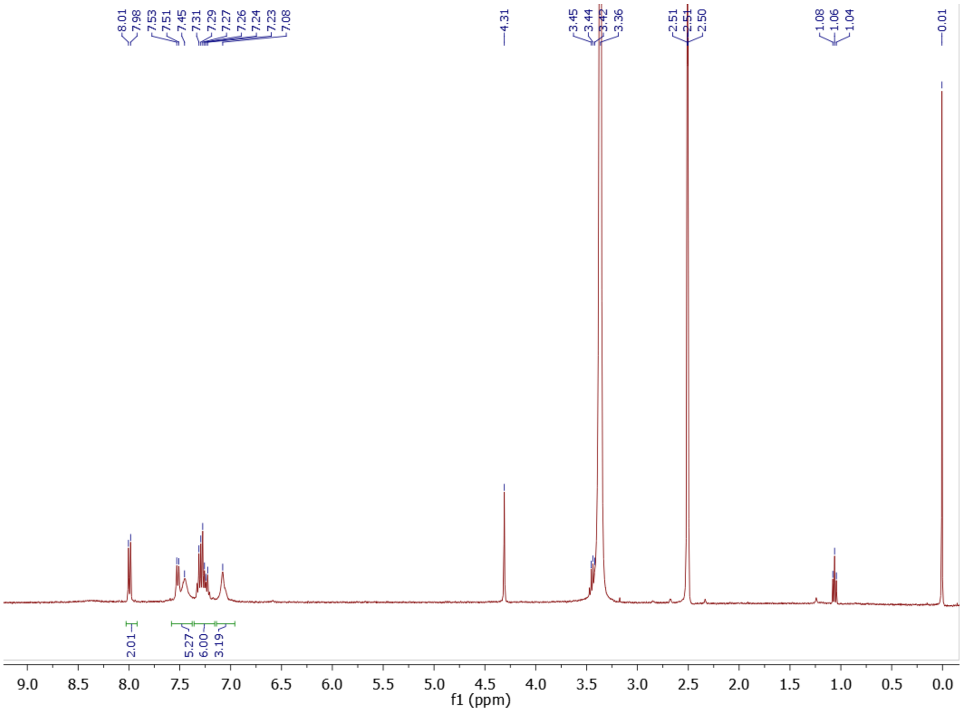

Figure S17: $^{1}$H NMR (400 MHz) spectrum of (4-(phenylethynylphenyl)biguanide trifluoroacetic salt **(3)** in DMSO-d6 at 298K

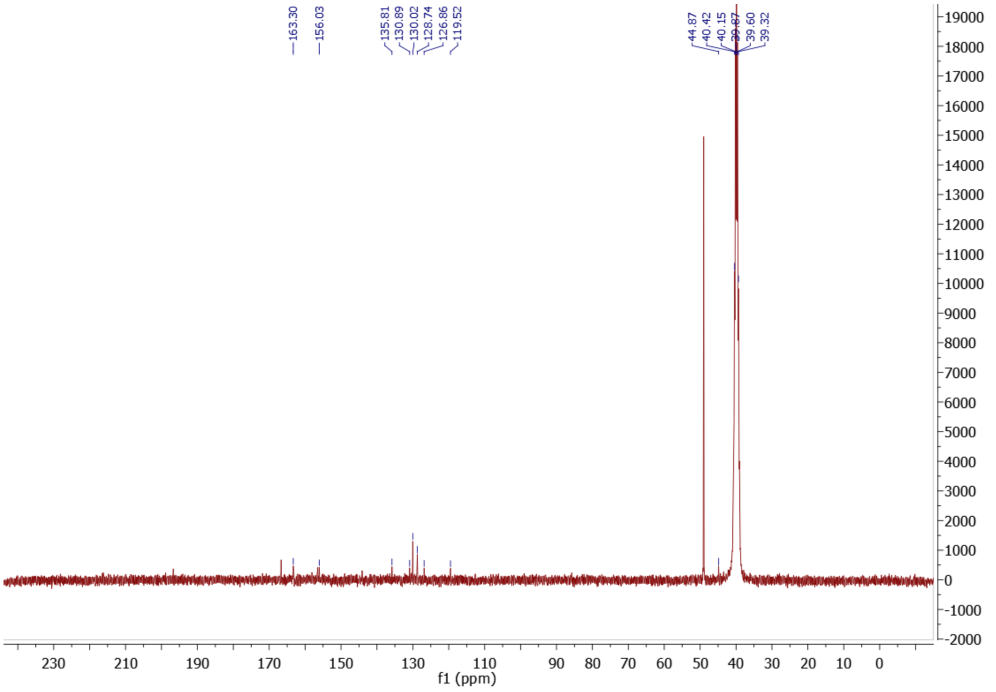

Figure S18: $^{13}$C NMR (70 MHz) spectrum of (4-(phenylethynylphenyl)biguanide trifluoroacetic salt **(3)** in DMSO-d6 at 298K

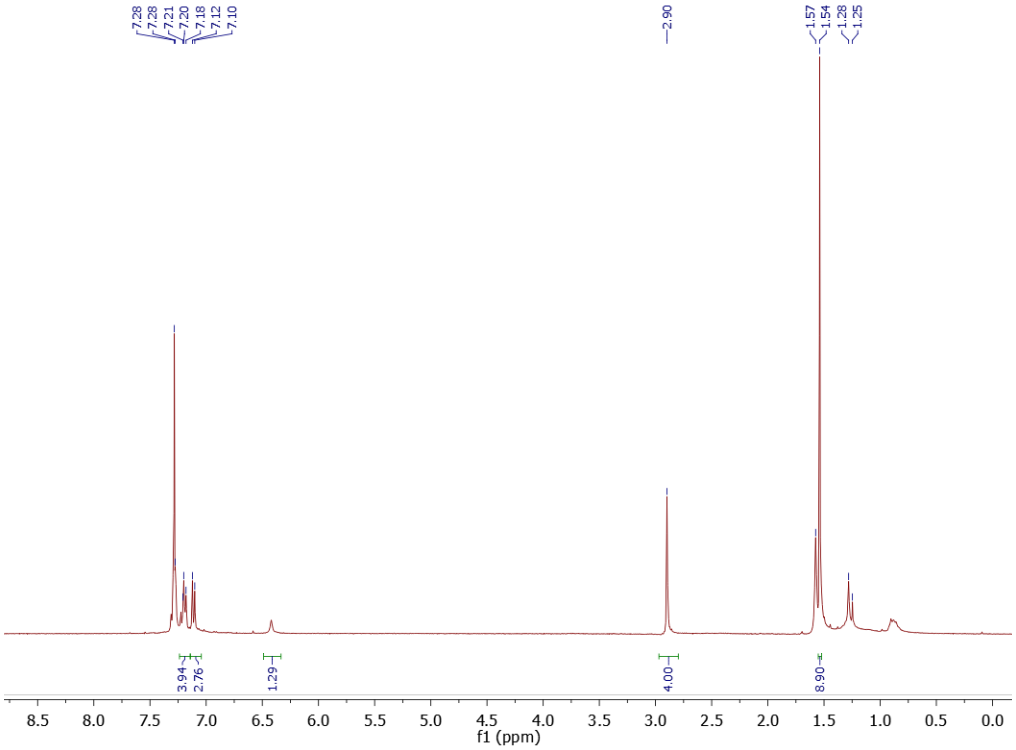

Figure S19: $^{1}$H NMR (400 MHz) spectrum of tert-butyl (4-phenethylphenyl)carbamate in $CDCl_3$ at 298K

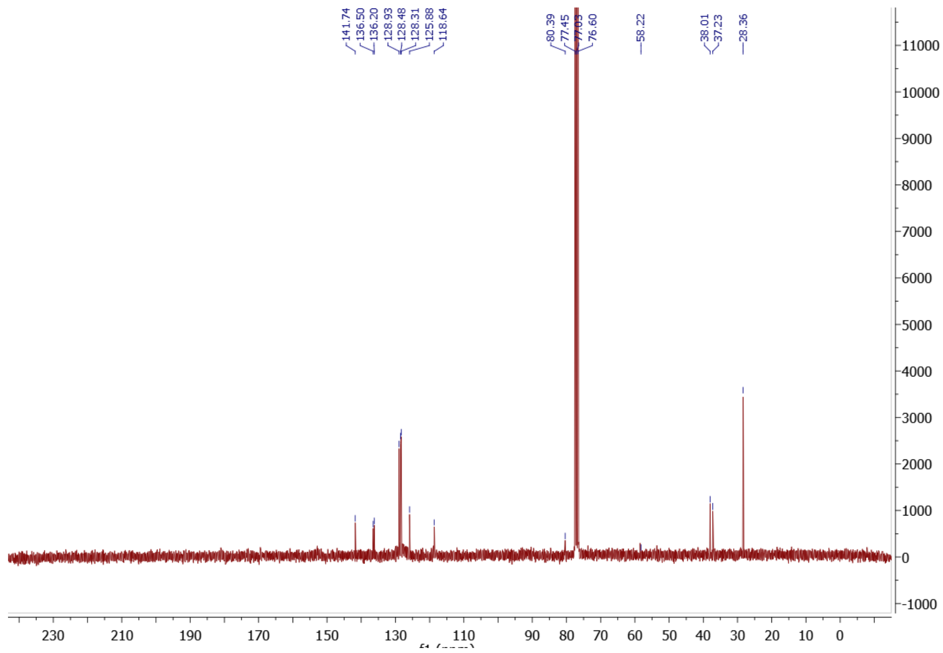

Figure S20: $^{13}$C NMR (70 MHz) spectrum of tert-butyl (4-phenethylphenyl)carbamate in $CDCl_3$ at 298K

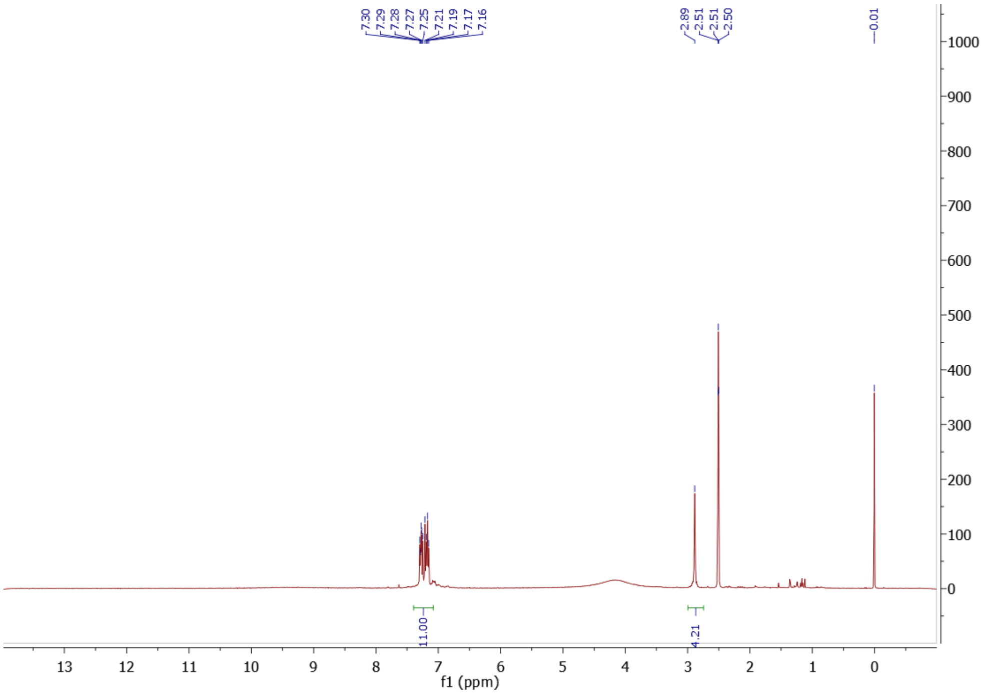

Figure S21: $^{1}H$ NMR (400 MHz) spectrum of 4-(phenylethylphenyl)aniline trifluoroacetic salt in DMSO-d6 at 298K

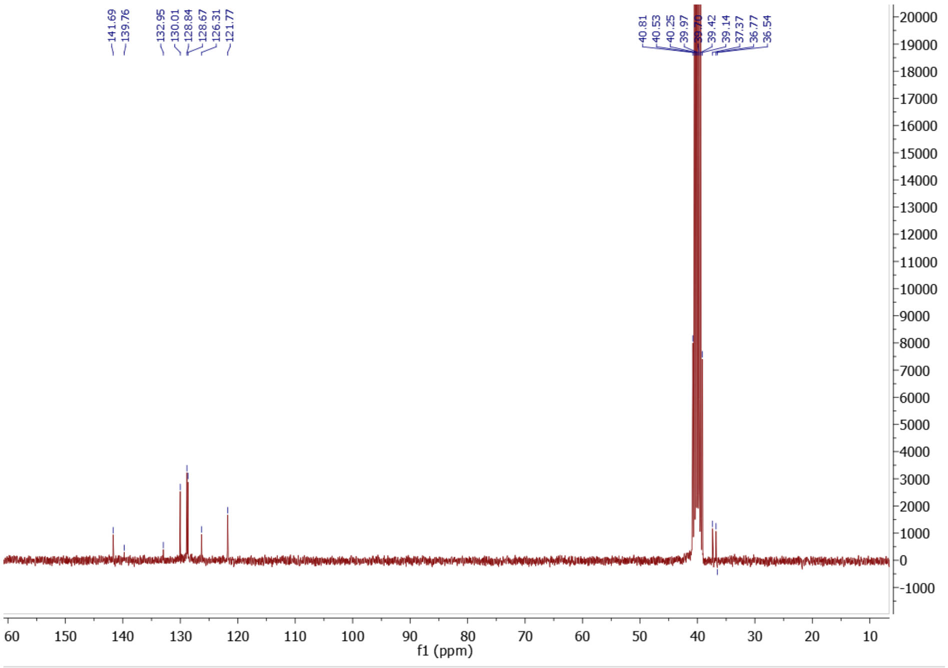

Figure S22: $^{13}C$ NMR (70 MHz) spectrum of 4-(phenylethylphenyl)aniline trifluoroacetic salt in DMSO-d6 at 298K

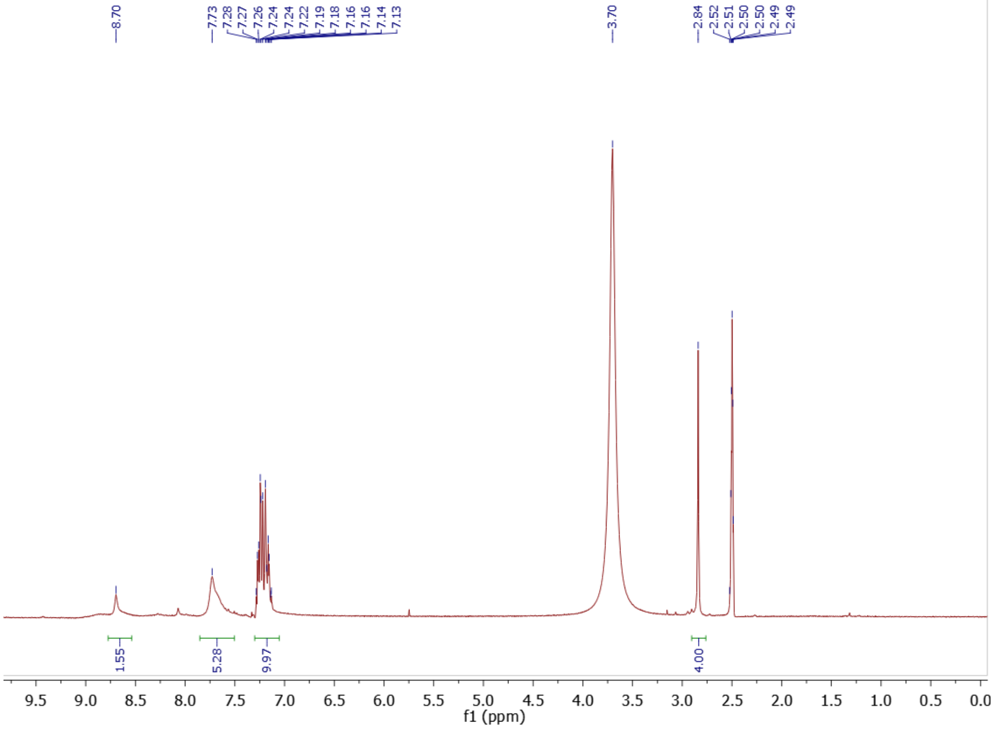

Figure S23: $^{1}$H NMR (400 MHz) spectrum of (4-(phenylethylphenyl)biguanide trifluoroacetic acid salt **(4)** in DMSO-d6 at 298K

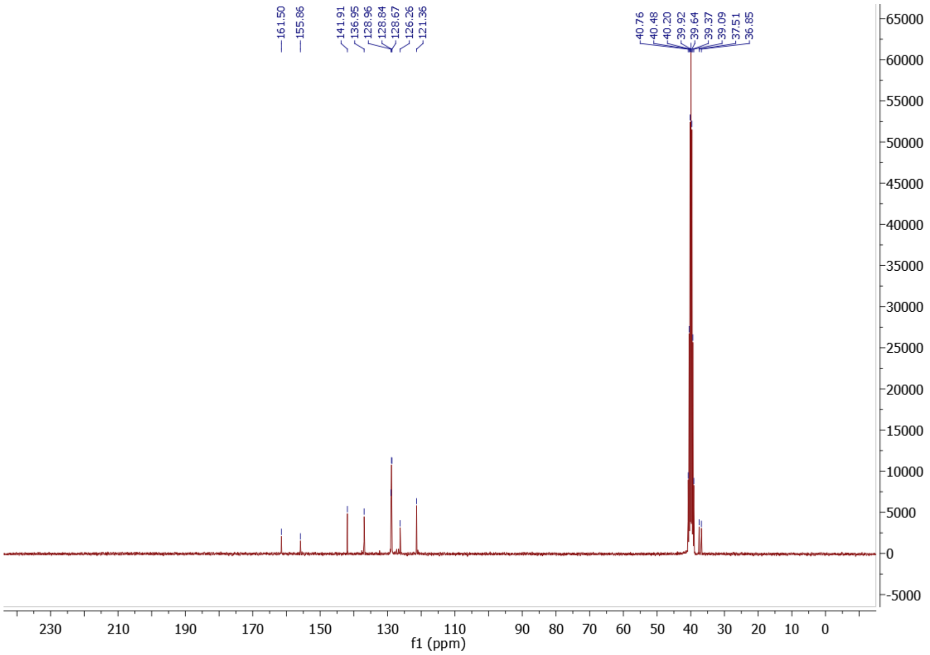

Figure S24: $^{13}$C NMR (70 MHz) spectrum of (4-(phenylethylphenyl)biguanide trifluoroacetic acid salt **(4)** in DMSO-d6 at 298K

### 2. Single crystal X-ray diffraction

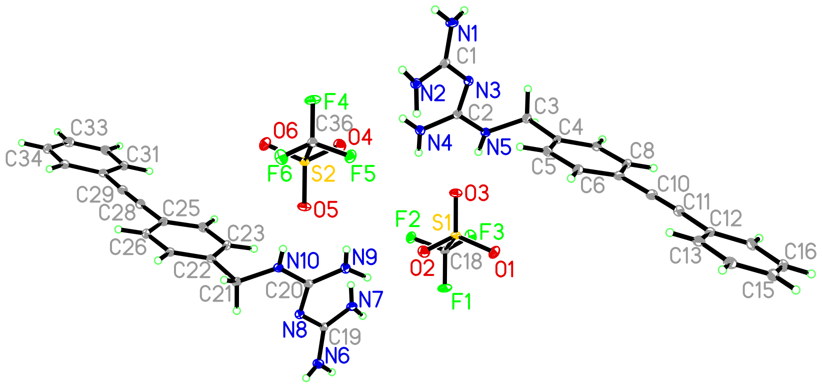

**Table 1 Crystal data and structure refinement for compound 1b**

| | |
|---|---|
| Identification code | 1b |
| Empirical formula | $C_{18}H_{18}F_3N_5O_3S$ |
| Formula weight | 441.43 |
| Temperature/K | 100 |
| Crystal system | orthorhombic |
| Space group | $Pca2_1$ |
| a/Å | 15.3543(10) |
| b/Å | 5.8490(4) |
| c/Å | 43.017(3) |
| α/° | 90 |
| β/° | 90 |
| γ/° | 90 |
| Volume/Å$^3$ | 3863.2(4) |
| Z | 8 |
| $\rho_{calc}$g/cm$^3$ | 1.518 |

| | |
|---|---|
| μ/mm$^{-1}$ | 1.321 |
| F(000) | 1824.0 |
| Crystal size/mm$^3$ | 0.34 × 0.14 × 0.11 |
| Radiation | GaKα (λ = 1.34139) |
| 2Θ range for data collection/° | 7.152 to 126.968 |
| Index ranges | -17 ≤ h ≤ 20, -7 ≤ k ≤ 7, -57 ≤ l ≤ 57 |
| Reflections collected | 81637 |
| Independent reflections | 9587 [$R_{int}$ = 0.0481, $R_{sigma}$ = 0.0246] |
| Data/restraints/parameters | 9587/1/599 |
| Goodness-of-fit on F$^2$ | 1.059 |
| Final R indexes [I>=2σ (I)] | $R_1$ = 0.0361, $wR_2$ = 0.0943 |
| Final R indexes [all data] | $R_1$ = 0.0372, $wR_2$ = 0.0952 |
| Largest diff. peak/hole / e Å$^{-3}$ | 0.34/-0.61 |
| Flack parameter | 0.22(2) |

**Table 2 Fractional Atomic Coordinates (×10$^4$) and Equivalent Isotropic Displacement Parameters (Å$^2$×10$^3$) for compound 1b. $U_{eq}$ is defined as 1/3 of of the trace of the orthogonalised $U_{IJ}$ tensor.**

| **Atom** | ***x*** | ***y*** | ***z*** | **U(eq)** |
|---|---|---|---|---|
| S1 | 4827.7(4) | 3450.5(10) | 4584.9(2) | 14.40(13) |
| F1 | 5669.4(11) | 30(3) | 4836.1(5) | 28.3(4) |
| F2 | 4285.2(11) | -12(3) | 4915.9(4) | 24.6(4) |
| F3 | 4793.7(13) | -931(3) | 4465.6(4) | 26.7(4) |
| O1 | 5538.7(14) | 3679(4) | 4364.1(5) | 24.8(4) |
| O2 | 4920.9(14) | 4715(4) | 4869.3(6) | 25.4(5) |
| O3 | 3969.4(14) | 3569(3) | 4443.9(5) | 21.5(4) |
| C18 | 4904.1(17) | 464(5) | 4706.7(6) | 16.3(5) |

| | | | | |
|---|---|---|---|---|
| S2 | 2468.6(4) | 11553.4(10) | 5398.3(2) | 14.62(13) |
| F4 | 1629.0(11) | 14971(3) | 5146.9(5) | 29.1(4) |
| F5 | 3011.6(11) | 15008(3) | 5066.3(4) | 25.6(4) |
| F6 | 2504.1(12) | 15936(3) | 5517.7(4) | 26.9(4) |
| O4 | 2375.5(14) | 10287(4) | 5114.7(5) | 25.7(5) |
| O5 | 3329.1(13) | 11434(3) | 5540.4(5) | 21.6(4) |
| O6 | 1762.5(14) | 11322(4) | 5617.7(5) | 24.3(4) |
| C36 | 2393.3(17) | 14538(5) | 5278.3(6) | 16.7(5) |
| N1 | 1171.2(17) | 12626(5) | 3977.1(6) | 22.8(5) |
| N2 | 2043.4(16) | 12865(4) | 4407.7(5) | 18.6(4) |
| N3 | 2239.3(14) | 9955(4) | 4026.8(5) | 16.3(4) |
| N4 | 2664.3(17) | 8290(4) | 4510.6(5) | 17.8(5) |
| N5 | 3216.6(15) | 7013(4) | 4051.8(5) | 16.0(4) |
| C1 | 1824.2(17) | 11771(5) | 4144.3(6) | 15.5(5) |
| C2 | 2699.6(18) | 8511(4) | 4200.4(6) | 14.6(5) |
| C3 | 3301.7(17) | 6869(5) | 3715.1(6) | 15.6(5) |
| C4 | 4075.3(16) | 8180(4) | 3587.5(6) | 14.0(5) |
| C5 | 4328.5(17) | 10272(5) | 3714.3(6) | 16.7(5) |
| C6 | 5006.0(19) | 11517(5) | 3583.0(7) | 18.0(5) |
| C7 | 5437.2(17) | 10689(5) | 3316.7(6) | 16.9(5) |
| C8 | 5186.7(17) | 8563(5) | 3192.9(6) | 17.8(5) |
| C9 | 4516.8(17) | 7330(5) | 3327.6(6) | 15.8(5) |
| C10 | 6129.6(18) | 11993(5) | 3179.3(7) | 19.2(5) |
| C11 | 6698.0(19) | 13128(5) | 3072.8(7) | 19.8(5) |
| C12 | 7387.1(17) | 14580(5) | 2961.1(6) | 17.1(5) |
| C13 | 7513.9(18) | 16731(5) | 3099.3(7) | 19.4(5) |
| C14 | 8178(2) | 18147(5) | 2995.8(7) | 22.6(6) |

| | | | | |
|---|---|---|---|---|
| C15 | 8717.3(19) | 17456(6) | 2752.5(7) | 24.2(6) |
| C16 | 8599(2) | 15318(6) | 2615.8(7) | 23.9(6) |
| C17 | 7938.3(19) | 13894(5) | 2717.7(6) | 21.0(5) |
| N6 | 6127.4(17) | 2377(5) | 6005.8(6) | 22.5(5) |
| N7 | 5259.0(16) | 2150(4) | 5576.5(6) | 18.0(4) |
| N8 | 5060.3(14) | 5060(4) | 5957.6(6) | 17.2(4) |
| N9 | 4634.4(17) | 6723(4) | 5473.2(5) | 19.0(5) |
| N10 | 4087.6(15) | 8006(4) | 5933.1(5) | 15.9(4) |
| C19 | 5470.0(17) | 3242(5) | 5838.9(6) | 14.9(5) |
| C20 | 4603.1(17) | 6512(4) | 5785.2(6) | 14.3(5) |
| C21 | 4000.9(17) | 8144(4) | 6267.3(6) | 15.6(5) |
| C22 | 3226.8(17) | 6830(4) | 6393.7(6) | 14.4(5) |
| C23 | 2971.7(17) | 4735(5) | 6265.8(6) | 17.0(5) |
| C24 | 2301.0(19) | 3480(5) | 6396.6(6) | 18.0(5) |
| C25 | 1859.2(17) | 4292(5) | 6660.0(6) | 16.9(5) |
| C26 | 2099.5(18) | 6414(5) | 6786.2(6) | 16.7(5) |
| C27 | 2774.9(17) | 7666(5) | 6652.7(6) | 15.5(5) |
| C28 | 1174.0(18) | 2953(5) | 6795.6(6) | 18.5(5) |
| C29 | 608.4(19) | 1764(5) | 6902.1(6) | 18.1(5) |
| C30 | -56.4(17) | 281(5) | 7017.5(7) | 17.1(5) |
| C31 | -203.5(18) | -1842(5) | 6874.0(7) | 18.4(5) |
| C32 | -846(2) | -3285(5) | 6984.6(7) | 22.1(6) |
| C33 | -1354.7(19) | -2639(6) | 7237.8(7) | 25.8(6) |
| C34 | -1219.7(19) | -556(6) | 7380.5(7) | 23.9(6) |
| C35 | -579.4(19) | 912(5) | 7273.4(7) | 20.3(5) |

**Table 3 Anisotropic Displacement Parameters (Å$^2$×10$^3$) for compound 1b. The Anisotropic displacement factor exponent takes the form: $-2\pi^2[h^2a^{*2}U_{11}+2hka^*b^*U_{12}+...]$.**

| Atom | $U_{11}$ | $U_{22}$ | $U_{33}$ | $U_{23}$ | $U_{13}$ | $U_{12}$ |
|---|---|---|---|---|---|---|
| S1 | 11.0(3) | 15.5(3) | 16.7(3) | 0.5(2) | -0.9(2) | 0.2(2) |
| F1 | 16.9(8) | 28.8(9) | 39.1(10) | 7.7(8) | -7.3(7) | 7.9(8) |
| F2 | 22.2(8) | 28.5(9) | 23.1(8) | 10.1(7) | 7.2(6) | 2.3(8) |
| F3 | 36.8(10) | 17.5(8) | 25.9(9) | -3.2(7) | 1.1(7) | -0.5(7) |
| O1 | 20.2(10) | 23.5(10) | 30.6(11) | 4.6(8) | 8.2(8) | -2.9(8) |
| O2 | 27.7(11) | 24.1(10) | 24.5(12) | -6.5(9) | -3.5(8) | 1.5(9) |
| O3 | 16.6(9) | 20.7(10) | 27.1(10) | 6.8(8) | -6.8(8) | -0.1(8) |
| C18 | 12.3(11) | 18.8(12) | 17.9(12) | 2.5(10) | 0.4(9) | 0.8(9) |
| S2 | 11.3(3) | 16.1(3) | 16.5(3) | 0.6(2) | -1.0(2) | -0.1(2) |
| F4 | 16.8(7) | 30.2(9) | 40.2(11) | 6.9(8) | -6.8(7) | 6.2(8) |
| F5 | 22.2(8) | 29.6(9) | 24.9(8) | 10.3(7) | 7.7(7) | 0.8(8) |
| F6 | 37.5(11) | 18.5(8) | 24.6(9) | -2.7(7) | 0.2(7) | -0.2(7) |
| O4 | 29.1(11) | 23.9(11) | 24.1(11) | -7.4(9) | -3.6(8) | 1.8(9) |
| O5 | 16.1(9) | 23.1(10) | 25.5(10) | 6.5(8) | -7.7(8) | -0.5(7) |
| O6 | 20.6(10) | 24.1(10) | 28.1(10) | 3.6(8) | 8.6(8) | -2.8(8) |
| C36 | 14.5(11) | 20.4(13) | 15.3(12) | 2.4(10) | 0.3(9) | 0.7(9) |
| N1 | 18.2(11) | 25.8(12) | 24.4(12) | -7.6(10) | -5.4(9) | 8.5(10) |
| N2 | 16.0(11) | 19.3(11) | 20.4(11) | -4.6(9) | -1.4(8) | 2.5(9) |
| N3 | 15.7(10) | 18.2(11) | 15.1(10) | -0.7(8) | 0.3(8) | 1.8(10) |
| N4 | 18.8(11) | 19.6(11) | 15.0(11) | 2.3(9) | -0.4(9) | 2.6(10) |
| N5 | 14.3(10) | 17.7(10) | 16.2(10) | 2.8(8) | 0.4(8) | 3.1(8) |
| C1 | 11.1(11) | 16.2(11) | 19.2(12) | 1.6(9) | 2.0(9) | -1.5(9) |
| C2 | 11.2(11) | 15.5(12) | 17.1(11) | 0.5(9) | 0.2(9) | -3.4(9) |
| C3 | 14.4(11) | 16.6(11) | 15.7(11) | -1.9(9) | 1.4(9) | -0.9(9) |
| C4 | 11.9(11) | 15.6(11) | 14.4(11) | 1.4(9) | -1.3(8) | 0.8(9) |

| | | | | | | |
|---|---|---|---|---|---|---|
| C5 | 17.8(12) | 17.7(12) | 14.8(11) | -0.1(9) | -0.6(9) | -2.2(10) |
| C6 | 17.8(12) | 16.2(12) | 20.1(12) | 0.6(9) | -2.7(10) | -1.2(10) |
| C7 | 13.2(11) | 19.3(12) | 18.1(12) | 3.1(10) | -1.6(9) | -0.9(9) |
| C8 | 13.6(12) | 22.1(13) | 17.5(11) | -1(1) | 0.7(9) | 3(1) |
| C9 | 15.1(11) | 16.9(12) | 15.4(11) | -1.6(9) | -2.1(9) | 2.3(9) |
| C10 | 16.7(13) | 20.1(12) | 20.8(12) | 1(1) | -1.6(10) | -0.3(10) |
| C11 | 18.7(13) | 22.7(13) | 18.1(11) | 2.3(10) | 0.1(10) | -0.4(10) |
| C12 | 14.4(11) | 20.6(12) | 16.4(12) | 4.2(10) | -3.3(9) | -3.4(10) |
| C13 | 16.5(13) | 21.0(14) | 20.7(12) | 2.1(11) | -2.3(10) | 1(1) |
| C14 | 23.8(14) | 20.0(12) | 24.0(13) | 2.2(10) | -6.3(10) | -4.3(11) |
| C15 | 21.3(13) | 30.1(15) | 21.3(13) | 5.4(11) | -1.8(11) | -9.7(12) |
| C16 | 21.3(13) | 33.5(15) | 16.9(12) | 3.3(11) | 0.6(10) | -4.1(12) |
| C17 | 23.4(13) | 23.0(13) | 16.7(11) | 2.1(10) | -1.3(10) | -3.5(11) |
| N6 | 19.5(11) | 25.2(12) | 22.9(12) | -6.8(10) | -4.4(9) | 8.4(10) |
| N7 | 13.4(11) | 19.0(11) | 21.7(11) | -2.7(9) | -2.6(9) | 1.7(9) |
| N8 | 15.1(10) | 21.0(12) | 15.5(11) | 0.3(8) | 0.1(7) | 2.8(10) |
| N9 | 18.8(11) | 22.2(12) | 16.1(11) | 2.1(9) | 0.7(8) | 1.2(10) |
| N10 | 14.7(10) | 17.9(10) | 15(1) | 3.7(8) | -1.5(8) | 1.7(9) |
| C19 | 12.0(11) | 18.6(12) | 13.9(11) | 1.2(9) | 3.0(9) | -2.1(9) |
| C20 | 11.0(11) | 16.6(12) | 15.2(11) | 0.1(9) | 1.3(9) | -3.8(9) |
| C21 | 13.0(11) | 16.9(11) | 17.0(11) | -1.1(9) | 1.4(9) | -1.6(9) |
| C22 | 12.5(11) | 15.5(11) | 15.1(11) | 1.8(9) | -2.1(9) | 1.1(9) |
| C23 | 16.0(12) | 19.0(12) | 16.0(11) | -1.2(9) | 0.5(9) | 1.1(10) |
| C24 | 18.6(12) | 17.1(13) | 18.3(12) | -1.2(9) | -1.7(10) | -2.3(9) |
| C25 | 13.9(11) | 20.6(12) | 16.4(11) | 4.5(9) | -1.6(9) | -2.7(9) |
| C26 | 14.9(12) | 20.6(13) | 14.6(11) | 0.0(9) | 2.0(9) | 0.1(10) |
| C27 | 13.9(11) | 16.6(11) | 16.0(11) | -0.7(9) | -2.0(9) | -0.7(9) |

| | | | | | | |
|---|---|---|---|---|---|---|
| C28 | 17.1(12) | 21.2(13) | 17.1(12) | 1.9(10) | -0.4(10) | -2.6(10) |
| C29 | 16.1(12) | 20.4(13) | 17.8(11) | 0.3(9) | -2.8(10) | -1.6(10) |
| C30 | 14.2(11) | 18.9(12) | 18.2(12) | 2.1(10) | -3.0(9) | 0(1) |
| C31 | 17.7(12) | 19.8(13) | 17.7(12) | -0.4(10) | -1.5(9) | -1.1(10) |
| C32 | 22.7(14) | 21.1(13) | 22.5(13) | 1.7(10) | -6.1(11) | -5.2(11) |
| C33 | 18.4(13) | 33.8(15) | 25.3(14) | 10.4(12) | -1.4(11) | -8.6(12) |
| C34 | 17.9(12) | 34.2(15) | 19.5(12) | 4.5(12) | 1.8(10) | 2.3(11) |
| C35 | 18.8(12) | 22.3(13) | 19.8(12) | -0.7(10) | -4.3(10) | 0.7(11) |

**Table 4 Bond Lengths for compound 1b.**

| Atom | Atom | Length/Å | Atom | Atom | Length/Å |
|---|---|---|---|---|---|
| S1 | O1 | 1.453(2) | C11 | C12 | 1.439(4) |
| S1 | O2 | 1.436(2) | C12 | C13 | 1.405(4) |
| S1 | O3 | 1.452(2) | C12 | C17 | 1.405(4) |
| S1 | C18 | 1.828(3) | C13 | C14 | 1.387(4) |
| F1 | C18 | 1.325(3) | C14 | C15 | 1.394(4) |
| F2 | C18 | 1.338(3) | C15 | C16 | 1.394(5) |
| F3 | C18 | 1.331(3) | C16 | C17 | 1.384(4) |
| S2 | O4 | 1.434(2) | N6 | C19 | 1.338(4) |
| S2 | O5 | 1.457(2) | N7 | C19 | 1.337(4) |
| S2 | O6 | 1.444(2) | N8 | C19 | 1.337(4) |
| S2 | C36 | 1.824(3) | N8 | C20 | 1.328(3) |
| F4 | C36 | 1.327(3) | N9 | C20 | 1.349(3) |
| F5 | C36 | 1.345(3) | N10 | C20 | 1.340(3) |
| F6 | C36 | 1.326(3) | N10 | C21 | 1.446(3) |
| N1 | C1 | 1.331(4) | C21 | C22 | 1.517(4) |

| | | | | | |
|---|---|---|---|---|---|
| N2 | C1 | 1.344(4) | C22 | C23 | 1.399(4) |
| N3 | C1 | 1.338(3) | C22 | C27 | 1.401(4) |
| N3 | C2 | 1.331(4) | C23 | C24 | 1.384(4) |
| N4 | C2 | 1.342(4) | C24 | C25 | 1.404(4) |
| N5 | C2 | 1.344(3) | C25 | C26 | 1.404(4) |
| N5 | C3 | 1.457(3) | C25 | C28 | 1.435(4) |
| C3 | C4 | 1.517(4) | C26 | C27 | 1.393(4) |
| C4 | C5 | 1.395(4) | C28 | C29 | 1.203(4) |
| C4 | C9 | 1.399(4) | C29 | C30 | 1.429(4) |
| C5 | C6 | 1.390(4) | C30 | C31 | 1.405(4) |
| C6 | C7 | 1.409(4) | C30 | C35 | 1.412(4) |
| C7 | C8 | 1.406(4) | C31 | C32 | 1.382(4) |
| C7 | C10 | 1.436(4) | C32 | C33 | 1.393(5) |
| C8 | C9 | 1.384(4) | C33 | C34 | 1.380(5) |
| C10 | C11 | 1.188(4) | C34 | C35 | 1.384(4) |

**Table 5 Bond Angles for compound 1b.**

| Atom | Atom | Atom | Angle/° | Atom | Atom | Atom | Angle/° |
|---|---|---|---|---|---|---|---|
| O1 | S1 | C18 | 103.12(12) | C9 | C8 | C7 | 120.4(2) |
| O2 | S1 | O1 | 115.75(14) | C8 | C9 | C4 | 120.6(2) |
| O2 | S1 | O3 | 114.93(13) | C11 | C10 | C7 | 177.8(3) |
| O2 | S1 | C18 | 103.98(14) | C10 | C11 | C12 | 176.5(3) |
| O3 | S1 | O1 | 113.83(13) | C13 | C12 | C11 | 119.3(3) |
| O3 | S1 | C18 | 102.92(12) | C17 | C12 | C11 | 121.6(3) |
| F1 | C18 | S1 | 111.14(19) | C17 | C12 | C13 | 119.2(3) |
| F1 | C18 | F2 | 107.9(2) | C14 | C13 | C12 | 120.0(3) |

| | | | | | | | |
|---|---|---|---|---|---|---|---|
| F1 | C18 | F3 | 108.8(2) | C13 | C14 | C15 | 120.3(3) |
| F2 | C18 | S1 | 110.24(19) | C16 | C15 | C14 | 120.0(3) |
| F3 | C18 | S1 | 110.76(19) | C17 | C16 | C15 | 120.1(3) |
| F3 | C18 | F2 | 107.8(2) | C16 | C17 | C12 | 120.4(3) |
| O4 | S2 | O5 | 114.98(13) | C20 | N8 | C19 | 123.0(3) |
| O4 | S2 | O6 | 115.65(13) | C20 | N10 | C21 | 124.2(2) |
| O4 | S2 | C36 | 104.31(14) | N7 | C19 | N6 | 117.1(3) |
| O5 | S2 | C36 | 102.83(12) | N8 | C19 | N6 | 116.8(3) |
| O6 | S2 | O5 | 113.70(13) | N8 | C19 | N7 | 126.0(3) |
| O6 | S2 | C36 | 103.12(12) | N8 | C20 | N9 | 126.5(3) |
| F4 | C36 | S2 | 111.1(2) | N8 | C20 | N10 | 117.7(2) |
| F4 | C36 | F5 | 107.2(2) | N10 | C20 | N9 | 115.7(2) |
| F5 | C36 | S2 | 110.04(19) | N10 | C21 | C22 | 113.6(2) |
| F6 | C36 | S2 | 111.24(19) | C23 | C22 | C21 | 121.5(2) |
| F6 | C36 | F4 | 109.1(2) | C23 | C22 | C27 | 118.7(2) |
| F6 | C36 | F5 | 108.1(2) | C27 | C22 | C21 | 119.7(2) |
| C2 | N3 | C1 | 123.0(2) | C24 | C23 | C22 | 120.8(2) |
| C2 | N5 | C3 | 124.3(2) | C23 | C24 | C25 | 120.6(3) |
| N1 | C1 | N2 | 117.7(3) | C24 | C25 | C26 | 119.0(2) |
| N1 | C1 | N3 | 116.9(3) | C24 | C25 | C28 | 119.9(3) |
| N3 | C1 | N2 | 125.3(3) | C26 | C25 | C28 | 121.2(3) |
| N3 | C2 | N4 | 126.7(3) | C27 | C26 | C25 | 120.1(2) |
| N3 | C2 | N5 | 117.4(2) | C26 | C27 | C22 | 120.8(2) |
| N4 | C2 | N5 | 115.7(3) | C29 | C28 | C25 | 177.5(3) |
| N5 | C3 | C4 | 113.6(2) | C28 | C29 | C30 | 177.4(3) |
| C5 | C4 | C3 | 121.4(2) | C31 | C30 | C29 | 119.9(3) |
| C5 | C4 | C9 | 119.3(2) | C31 | C30 | C35 | 118.8(3) |

| | | | | | | | |
|---|---|---|---|---|---|---|---|
| C9 | C4 | C3 | 119.3(2) | C35 | C30 | C29 | 121.2(3) |
| C6 | C5 | C4 | 120.6(2) | C32 | C31 | C30 | 120.2(3) |
| C5 | C6 | C7 | 120.1(3) | C31 | C32 | C33 | 120.3(3) |
| C6 | C7 | C10 | 120.0(3) | C34 | C33 | C32 | 120.2(3) |
| C8 | C7 | C6 | 118.9(2) | C33 | C34 | C35 | 120.4(3) |
| C8 | C7 | C10 | 121.1(3) | C34 | C35 | C30 | 120.1(3) |

**Table 6 Hydrogen Bonds for compound 1b.**

| D | H | A | d(D-H)/Å | d(H-A)/Å | d(D-A)/Å | D-H-A/° |
|---|---|---|---|---|---|---|
| N4 | H4A | O4 | 0.78(4) | 2.35(4) | 2.883(3) | 127(3) |
| N4 | H4A | N2 | 0.78(4) | 2.40(4) | 2.875(4) | 121(3) |
| N9 | H9A | O2 | 0.93(4) | 2.17(4) | 2.885(3) | 133(3) |
| N9 | H9A | N7 | 0.93(4) | 2.34(4) | 2.876(4) | 116(3) |
| N4 | H4B | F5[1] | 0.81(4) | 2.52(4) | 3.112(3) | 131(3) |
| N9 | H9B | F2[2] | 0.80(5) | 2.48(5) | 3.111(3) | 136(4) |
| N2 | H2A | O3[2] | 0.95(4) | 2.17(4) | 2.990(3) | 143(3) |
| N7 | H7A | O5[1] | 0.74(4) | 2.36(4) | 2.997(3) | 146(4) |
| N1 | H1A | O1[3] | 0.87(4) | 2.07(4) | 2.896(3) | 159(3) |
| N6 | H6A | O6[4] | 0.82(4) | 2.14(4) | 2.902(3) | 156(3) |
| N2 | H2B | O1[3] | 0.85(4) | 2.34(4) | 3.075(3) | 145(4) |
| N7 | H7B | O6[4] | 0.82(4) | 2.37(4) | 3.080(3) | 146(3) |
| N5 | H5 | O3 | 0.81(4) | 2.07(4) | 2.870(3) | 170(4) |
| N10 | H10 | O5 | 0.76(5) | 2.13(5) | 2.868(3) | 165(5) |

[1]+X,-1+Y,+Z; [2]+X,1+Y,+Z; [3]-1/2+X,2-Y,+Z; [4]1/2+X,1-Y,+Z

**Table 7 Torsion Angles for compound 1b.**

| A | B | C | D | Angle/° | A | B | C | D | Angle/° |
|---|---|---|---|---|---|---|---|---|---|
| O1 | S1 | C18 | F1 | 59.9(2) | C10 | C7 | C8 | C9 | -179.8(2) |
| O1 | S1 | C18 | F2 | 179.48(19) | C11 | C12 | C13 | C14 | 179.6(3) |
| O1 | S1 | C18 | F3 | -61.3(2) | C11 | C12 | C17 | C16 | -179.6(3) |
| O2 | S1 | C18 | F1 | -61.3(2) | C12 | C13 | C14 | C15 | 0.6(4) |
| O2 | S1 | C18 | F2 | 58.3(2) | C13 | C12 | C17 | C16 | 0.2(4) |
| O2 | S1 | C18 | F3 | 177.57(18) | C13 | C14 | C15 | C16 | -1.1(4) |
| O3 | S1 | C18 | F1 | 178.5(2) | C14 | C15 | C16 | C17 | 1.1(5) |
| O3 | S1 | C18 | F2 | -61.9(2) | C15 | C16 | C17 | C12 | -0.6(4) |
| O3 | S1 | C18 | F3 | 57.4(2) | C17 | C12 | C13 | C14 | -0.2(4) |
| O4 | S2 | C36 | F4 | 60.8(2) | N10 | C21 | C22 | C23 | 37.1(3) |
| O4 | S2 | C36 | F5 | -57.8(2) | N10 | C21 | C22 | C27 | -145.7(2) |
| O4 | S2 | C36 | F6 | -177.50(18) | C19 | N8 | C20 | N9 | -16.0(4) |
| O5 | S2 | C36 | F4 | -178.83(19) | C19 | N8 | C20 | N10 | 167.5(2) |
| O5 | S2 | C36 | F5 | 62.6(2) | C20 | N8 | C19 | N6 | 158.1(3) |
| O5 | S2 | C36 | F6 | -57.2(2) | C20 | N8 | C19 | N7 | -25.1(4) |
| O6 | S2 | C36 | F4 | -60.4(2) | C20 | N10 | C21 | C22 | -95.9(3) |
| O6 | S2 | C36 | F5 | -178.96(19) | C21 | N10 | C20 | N8 | 0.3(4) |
| O6 | S2 | C36 | F6 | 61.3(2) | C21 | N10 | C20 | N9 | -176.5(2) |
| N5 | C3 | C4 | C5 | -37.2(3) | C21 | C22 | C23 | C24 | 175.6(2) |
| N5 | C3 | C4 | C9 | 146.0(2) | C21 | C22 | C27 | C26 | -175.6(2) |
| C1 | N3 | C2 | N4 | 16.6(4) | C22 | C23 | C24 | C25 | 0.3(4) |
| C1 | N3 | C2 | N5 | -167.9(2) | C23 | C22 | C27 | C26 | 1.7(4) |

| | | | | | | | | | |
|---|---|---|---|---|---|---|---|---|---|
| C2 | N3 | C1 | N1 | -158.3(3) | C23 | C24 | C25 | C26 | 1.0(4) |
| C2 | N3 | C1 | N2 | 25.4(4) | C23 | C24 | C25 | C28 | -179.0(3) |
| C2 | N5 | C3 | C4 | 96.0(3) | C24 | C25 | C26 | C27 | -0.9(4) |
| C3 | N5 | C2 | N3 | -0.2(4) | C25 | C26 | C27 | C22 | -0.5(4) |
| C3 | N5 | C2 | N4 | 175.9(2) | C27 | C22 | C23 | C24 | -1.6(4) |
| C3 | C4 | C5 | C6 | -176.1(3) | C28 | C25 | C26 | C27 | 179.1(3) |
| C3 | C4 | C9 | C8 | 175.6(2) | C29 | C30 | C31 | C32 | 180.0(3) |
| C4 | C5 | C6 | C7 | 0.9(4) | C29 | C30 | C35 | C34 | 179.9(3) |
| C5 | C4 | C9 | C8 | -1.3(4) | C30 | C31 | C32 | C33 | 0.4(4) |
| C5 | C6 | C7 | C8 | -1.8(4) | C31 | C30 | C35 | C34 | 0.5(4) |
| C5 | C6 | C7 | C10 | 179.2(3) | C31 | C32 | C33 | C34 | -0.1(4) |
| C6 | C7 | C8 | C9 | 1.2(4) | C32 | C33 | C34 | C35 | 0.1(4) |
| C7 | C8 | C9 | C4 | 0.3(4) | C33 | C34 | C35 | C30 | -0.3(4) |
| C9 | C4 | C5 | C6 | 0.6(4) | C35 | C30 | C31 | C32 | -0.5(4) |

**Table 8 Hydrogen Atom Coordinates (Å×10$^4$) and Isotropic Displacement Parameters (Å$^2$×10$^3$) for compound 1b.**

| Atom | *x* | *y* | *z* | U(eq) |
|---|---|---|---|---|
| H3A | 2763 | 7468 | 3618 | 19 |
| H3B | 3359 | 5241 | 3655 | 19 |
| H5A | 4035 | 10851 | 3892 | 20 |
| H6 | 5179 | 12929 | 3673 | 22 |
| H8 | 5479 | 7971 | 3016 | 21 |
| H9 | 4355 | 5892 | 3243 | 19 |
| H13 | 7144 | 17217 | 3264 | 23 |
| H14 | 8266 | 19594 | 3091 | 27 |
| H15 | 9165 | 18442 | 2680 | 29 |

| | | | | |
|---|---|---|---|---|
| H16 | 8973 | 14837 | 2452 | 29 |
| H17 | 7857 | 12444 | 2622 | 25 |
| H21A | 3943 | 9770 | 6328 | 19 |
| H21B | 4539 | 7545 | 6364 | 19 |
| H23 | 3263 | 4166 | 6087 | 20 |
| H24 | 2138 | 2058 | 6307 | 22 |
| H26 | 1801 | 6997 | 6963 | 20 |
| H27 | 2931 | 9106 | 6738 | 19 |
| H31 | 139 | -2288 | 6700 | 22 |
| H32 | -941 | -4722 | 6887 | 27 |
| H33 | -1796 | -3636 | 7313 | 31 |
| H34 | -1568 | -127 | 7553 | 29 |
| H35 | -492 | 2347 | 7372 | 24 |
| H4A | 2330(20) | 9070(60) | 4594(9) | 11(8) |
| H9A | 4970(30) | 5730(70) | 5353(10) | 29(10) |
| H4B | 3040(20) | 7510(60) | 4590(8) | 11(7) |
| H9B | 4330(30) | 7700(80) | 5402(11) | 37(11) |
| H2A | 2600(30) | 12650(70) | 4495(9) | 23(9) |
| H7A | 4820(30) | 2370(70) | 5512(9) | 21(10) |
| H1A | 870(20) | 13750(60) | 4058(8) | 15(8) |
| H6A | 6390(20) | 1250(60) | 5945(8) | 14(8) |
| H2B | 1760(30) | 14040(70) | 4463(9) | 26(10) |
| H7B | 5510(30) | 950(70) | 5535(9) | 20(9) |
| H5 | 3490(20) | 6130(60) | 4160(8) | 16(8) |
| H10 | 3800(30) | 8800(80) | 5836(11) | 39(12) |
| H1B | 970(30) | 11830(80) | 3849(12) | 40(13) |
| H6B | 6340(30) | 3290(80) | 6131(12) | 39(12) |

**Experimental**

Single crystals of $C_{18}H_{18}F_3N_5O_3S$ were **[]**. A suitable crystal was selected and **[]** on a **Bruker Venture Metaljet** diffractometer. The crystal was kept at 100 K during data collection. Using Olex, the structure was solved with the XT structure solution program using Intrinsic Phasing and refined with the XL refinement package using Least Squares minimisation.

**Crystal structure determination of 1b**

**Crystal Data** for $C_{18}H_{18}F_3N_5O_3S$ ($M$ =441.43 g/mol): orthorhombic, space group Pca$2_1$ (no. 29), $a$ = 15.3543(10) Å, $b$ = 5.8490(4) Å, $c$ = 43.017(3) Å, $V$ = 3863.2(4) Å$^3$, $Z$ = 8, $T$ = 100 K, μ(GaKα) = 1.321 mm$^{-1}$, $Dcalc$ = 1.518 g/cm$^3$, 81637 reflections measured (7.152° ≤ 2Θ ≤ 126.968°), 9587 unique ($R_{int}$ = 0.0481, $R_{sigma}$ = 0.0246) which were used in all calculations. The final $R_1$ was 0.0361 (I > 2σ(I)) and $wR_2$ was 0.0952 (all data).

**Refinement model description**

Number of restraints - 1, number of constraints - unknown. Details:

1. Twinned data refinement
 Scales: 0.78(2) 0.22(2)
2. Fixed Uiso
 At 1.2 times of:
  All C(H) groups, All C(H,H) groups
3.a Secondary CH2 refined with riding coordinates:
 C3(H3A,H3B), C21(H21A,H21B)
3.b Aromatic/amide H refined with riding coordinates:
 C5(H5A), C6(H6), C8(H8), C9(H9), C13(H13), C14(H14), C15(H15), C16(H16),
 C17(H17), C23(H23), C24(H24), C26(H26), C27(H27), C31(H31), C32(H32),
C33(H33),
  C34(H34), C35(H35)

#### 3. Hemolytic activity

Red blood cells in Alsever's solution were centrifuged for 10 min at 300*g*, washed 3 times with PBS buffer, and resuspended in PBS at 2% v/v. To each well of a 96-well plate, 195 µL of red blood cell solution and 5 µL of biguanidium salt in DMSO were added, and the plate was incubated with light agitation for 1 h at 37 °C. The plate was then centrifuged for 10 min at 300*g*, and 50 µL of the supernatant solution of each well was transferred to another plate. Absorbance was measured at λ = 405 nm. Each measurement was performed in triplicate in three different experiments.

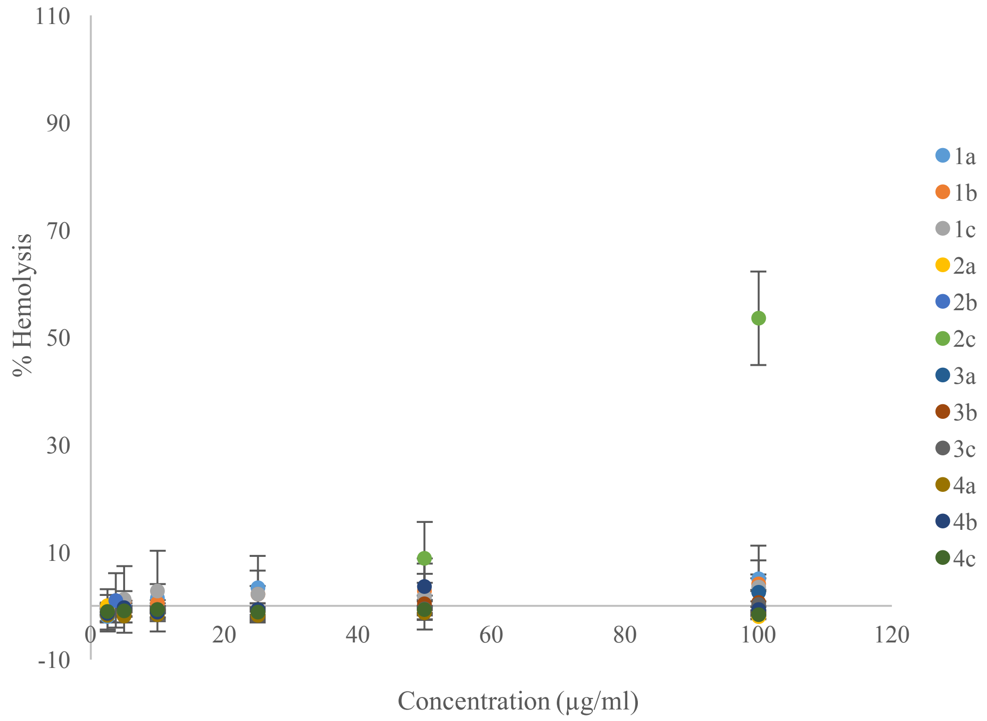

Figure S25: Hemolytic activity

Table 9: Minimal concentration for under 10% hemolysis

| | ***HC 10% (µg/ml)*** | ***HC 10% (µM)*** |
|---|---|---|
| ***1a*** | > 100 | > 175 |
| ***1b*** | > 100 | > 227 |
| ***1c*** | > 100 | > 305 |
| ***2a*** | > 100 | > 173 |
| ***2b*** | > 100 | > 225 |
| ***2c*** | 50 | 151 |
| ***3a*** | > 100 | > 179 |
| ***3b*** | > 100 | > 234 |
| ***3c*** | > 100 | > 319 |
| ***4a*** | > 100 | > 178 |
| ***4b*** | > 100 | > 232 |
| ***4c*** | > 100 | > 315 |

### 4. Measurement of the LogP

A 10 µM or 20 µM solution of **1b** was prepared in 5 ml of octanol and mixed with 5 ml distilled water at 25°C The solution was stirred and was left to settle for 24 hours before the absorbance of the octanol fraction was measured. The values obtained were fitted on a calibration curve prepared by measuring the absorbance of **1b** in octanol at various concentrations (2.5, 10, 15, 20 and 30 µM). The logP was calculated using the following formula:

$$logP = \log(\frac{[1b]_{oct}}{[1b]_{water}})$$

Where $[1b]_{oct}$ is the concentration of **1b** in octanol and $[1b]_{water}$ is the concentration of **1b** in water.

Table 10: Calibration curve and Log P values of compound **1b**

| Concentration of 1b (µM) | Absorbance (u.a) |
|---|---|
| 2.5 | 0.0908 |
| 10 | 0.3416 |
| 15 | 0.5334 |
| 20 | 0.6685 |
| 30 | 0.9611 |

| | |
|---|---|
| **Slope** | 0.0317 |
| **Intercept** | 0.0282 |

| | Absorbance (u.a) | [1b] in octanol (µM) | [1b] in water (µM) | logP | |
|---|---|---|---|---|---|
| Partition 1 (10 µM) | 0.2473 | 6.91 | 3.09 | 0.35 | 0.42 ± 0.10 |
| Partition 2 (20 µM) | 0.5086 | 15.15 | 4.85 | 0.50 | |

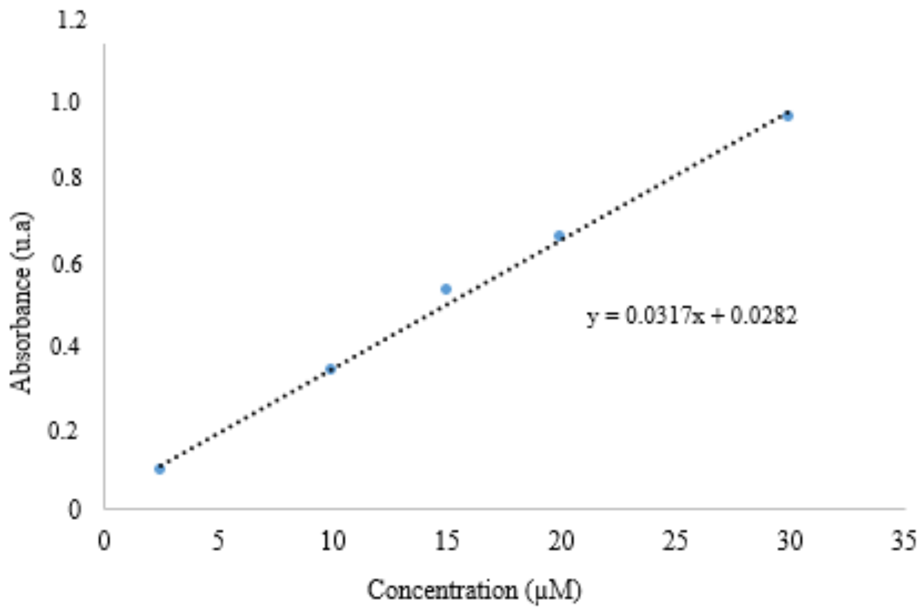

Figure S26: Calibration curve of 1b in octanol

### 5. U-Tube experiments

U-tube experiment was prepared by adding 800 µL of dichloromethane in a U-shaped tube as a representation of the hydrophobic lipid membrane. On the receiving end (*trans* side) of the tube was added 300 µL of distilled water while the other side (*cis* side) was filled with 300 µL of a 250 µM solution of the compound of interest at 25°C. At 48h and 72h, 100 µL aliquot of the *trans* side was diluted in 900 µL methanol and monitored by LCMS (292 m/z). Area under curve (AUC) was measured and fitted on a calibration curve.

Table 11: Calibration curve and U-tube experiment of compound **1b**

| **Concentration of 1b (µM)** | **Average AUC** |
|---|---|
| **0.125** | 6.11E+05 ± 7.71E+04 |
| **1.25** | 3.95E+06 ± 4.66E+05 |
| **12.5** | 2.05E+07 ± 1.41E+06 |
| **125** | 7.98E+07 ± 5.17E+06 |

| | |
|---|---|
| Slope | 6.02E+05 |
| Intercept | 5.32E+06 |

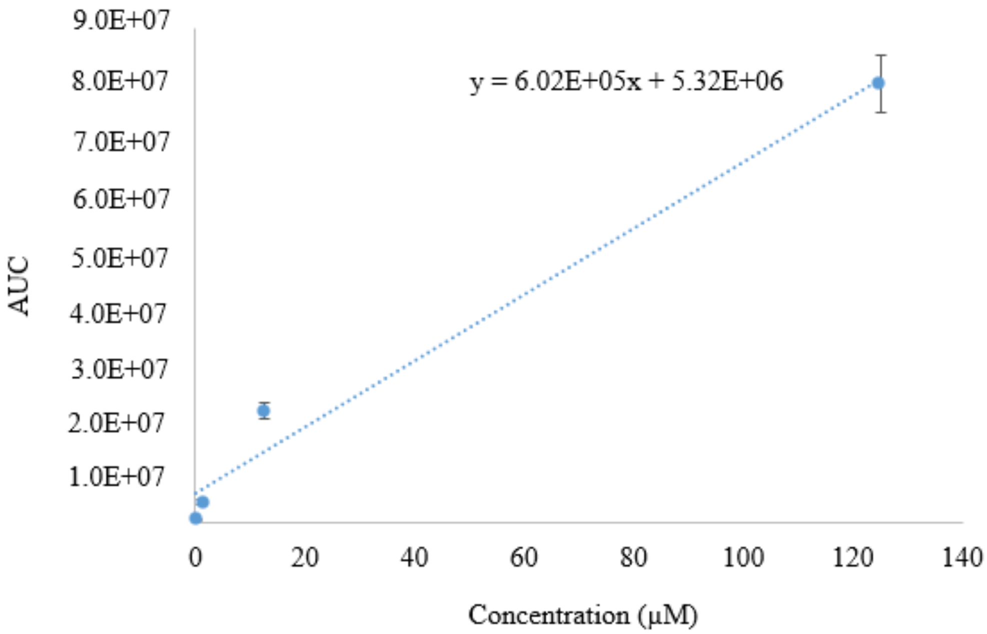

| | | AUC | Diluted concentration (µM) | Trans-side concentration (µM) | |
|---|---|---|---|---|---|
| **48h** | **1** | 1,07E+07 | 9,01 | 90,1 | 64,7 ± 22,6 |
| | **2** | 8,76E+06 | 5,71 | 57,1 | |
| | **3** | 8,14E+06 | 4,68 | 46,8 | |
| **72h** | **1** | 1,01E+07 | 7,86 | 78,6 | 92,3 ± 11,9 |
| | **2** | 1,13E+07 | 9,93 | 99,3 | |
| | **3** | 1,13E+07 | 9,91 | 99,1 | |

#### 6. Lucigenin assay

A 2 mL volume of a 25 mg/mL solution of EYPC in chloroform was slowly reduced *in vacuo* to form a thin film on the side of the flask. Then, 1 mL of a lucigenin solution (2 mM lucigenin, 10 mM $Na_2HPO_4$, 10 mM $NaH_2PO_4$, and 100 mM NaCl) was added, and the resulting suspension was subjected to 10 freeze/thaw cycles (1 cycle = 1 min at -20 °C and 1 min at 37 °C). The mixture was extruded onto a 100-nm polycarbonate membrane 21 times and passed through a Sephadex G-25 column to remove the extravesicular lucigenin. The eluent used for the column was a phosphate buffer with sodium chloride (10 mM $Na_2HPO_4$, 10 mM $NaH_2PO_4$, and 100 mM NaCl), and the resulting liposome solution was diluted to obtain a final concentration of 10 mM. To a quartz cuvette, 2.5 mL phosphate buffer with sodium nitrate (10 mM $Na_2HPO_4$, 10 mM $NaH_2PO_4$, and 100 mM $NaNO_3$) and 40 µL of liposome solution were added with light stirring ($\lambda_{ex.}$ = 372 nm, $\lambda_{em.}$ = 503 nm). At *t* = 50 s, a solution of the biguanidium in methanol was added to the cuvette to obtain a 50 mM final solution (50 mol% relative to the concentration of EYPC). At *t* = 300 s, Triton-X 10% v/v was added to lyse the liposomes. The fluorescence was monitored for 350 s.

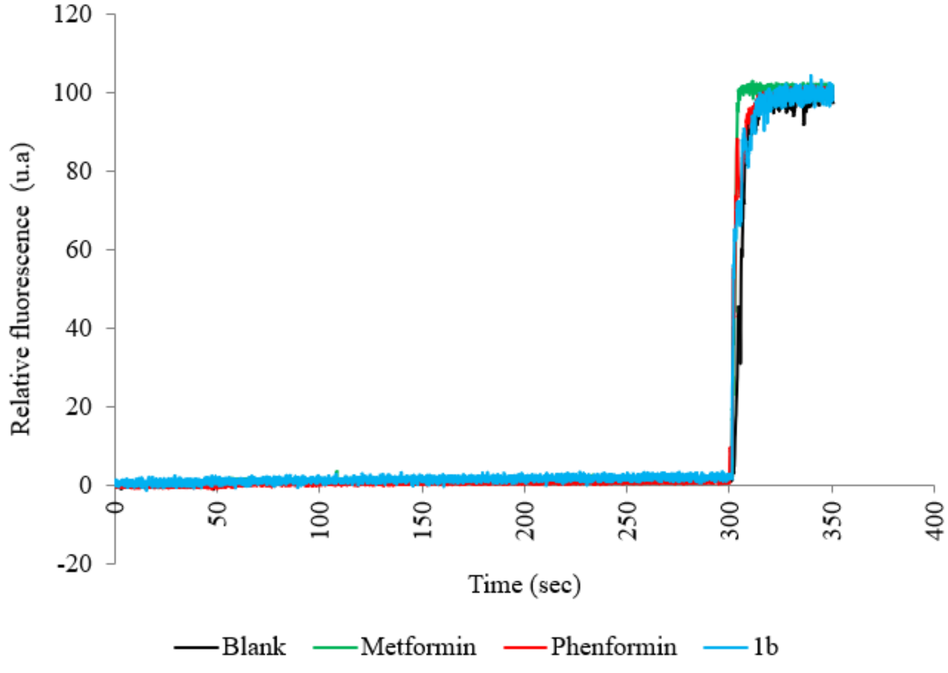

Figure S27: Chloride transport assay with lucigenin

### 7. HPTS assay

2 mL of a 25 mg/mL solution of EYPC in chloroform was slowly reduced *in vacuo* to form a thin film on the side of the flask. Then, 1 mL of a solution of the trisodium salt of 8-hydroxypyrene-1,3,6-trisulfonic acid (HPTS) (1 mM HPTS, 10 mM 4-(2-hydroxyethyl)-1-piperazineethanesulfonic acid (HEPES) salt, and 100 mM NaCl, adjusted to pH = 7.4) was added, and the resulting suspension was subjected to 10 freeze/thaw cycles (1 cycle = 1 min at -20 °C and 1 min at 37 °C). The mixture was extruded onto a 100-nm polycarbonate membrane 21 times and passed through a Sephadex G-25 column to remove the extravesicular HPTS. HEPES buffer (10 mM HEPES and 100 mM NaCl, adjusted to pH = 7.4) was used as the eluent for the column and the resulting liposome solution was diluted to obtain a final concentration of 10 mM. To a quartz cuvette, 1.9 mL of HEPES buffer and 25 μL of liposome solution were added with light stirring ($\lambda_{ex.}$ = 405/450 nm, $\lambda_{em.}$ = 510 nm). At *t* = 50 s, a solution biguanidium salt in methanol was added to obtain a 5 mM final concentration. At *t* = 300 s, NaOH was added to obtain a 5 mM final concentration, and, at *t* = 350 s, Triton-X 10% v/v was added to lyse the liposomes. The fluorescence was monitored for 600 s. As a control, we monitored the variation of HPTS fluorescence after the addition of a 100 μg/mL solution of biguanidium salts. No variation was observed.

### 8. Safranin O assay

2 mL of a 25 mg/mL solution of EYPC in chloroform was slowly reduced *in vacuo* to form a thin film on the side of the flask. Then, 1 mL of HEPES buffer (10 mM HEPES salt, 100 mM KCl, adjusted to pH = 7.4) was added, and the resulting suspension was subjected to 10 freeze/thaw cycles (1 cycle = 1 min at -20 °C and 1 min at 37 °C). The mixture was extruded on a 100 nm polycarbonate membrane 21 times, and the resulting liposome solution was diluted to obtain a 10 mM final concentration. To a quartz cuvette, 1.9 mL of HEPES buffer with sodium chloride (10 mM HEPES salt and 100 mM NaCl, adjusted to pH = 7.4) and 100 μL of liposome solution were added with light stirring. Safranin O dye

was added to a 60 nM final concentration ($\lambda_{ex.}$ = 522 nm, $\lambda_{em.}$ = 581 nm). At $t$ = 50 s, a solution of biguanidium salt in DMSO was added to obtain a 100 μg/mL final solution in the cuvette, and the fluorescence was monitored for 300 s. As a control, we monitored the variation in safranin O fluorescence in the solution by adding 100 μg/mL biguanidium salts. No change was observed.

#### 9. Mitochondrial permeation and accumulation

pMXs-3XHA-EGFP-OMP25 retroviral particles were produced in Phoenix cells and then incubated for 8 hours in KP4 cells. Then, positive cells were selected with blasticidine (Santa Cruz Biotechnology, sc-495389). At day 7 post-infection, ~20 million cells were treated for 3 hours with 15 μM metformin, or 15 μM compound **1b**, or DMSO as vehicle. After treatment, cells were washed twice with PBS and then scraped into 1 mL chilled KPBS (136 mM KCl, 10mM $KH_2PO_4$, pH 7.25) for mitochondrial isolation as described in the litterature[1], except for the incubation with anti-HA magnetic beads (Thermo Fisher Scientific, 88837), in which the supernatants were incubated with 30 μL of prewashed beads on a vertical-rotating mixer for 1 hour. pMXs-3XHA-EGFP-OMP25 was a gift from David Sabatini (Addgene plasmid # 83356; http://n2t.net/addgene:83356; RRID: Addgene_83356). A standard MTP target plate was used for MALDI-MS analysis (Bruker Daltonics, Billerica, MA). An organic matrix solution of *α*-cyano-4-hydroxycinnamic acid (*α*-CHCA) was best suited for the biguanide drugs and was prepared at a 7 mg/mL concentration in an equal ratio of acetonitrile (ACN) and $H_2O$. A 0.5 μL drop of the matrix solution was placed on the target plate for each biological solution analyzed and let to air dry. Then, for each sample, a 0.5 μL drop was pipetted on one of the dried matrix spots and let to air dry. For the MS experiment, an accumulation of 250 shots was obtained for each sample at *m/z* 0-1000 and repeated three times. External calibration was carried out in cubic enhanced mode using known matrix peaks and CsI to obtain five points of calibration over the considered mass range.

### 10. Full gel

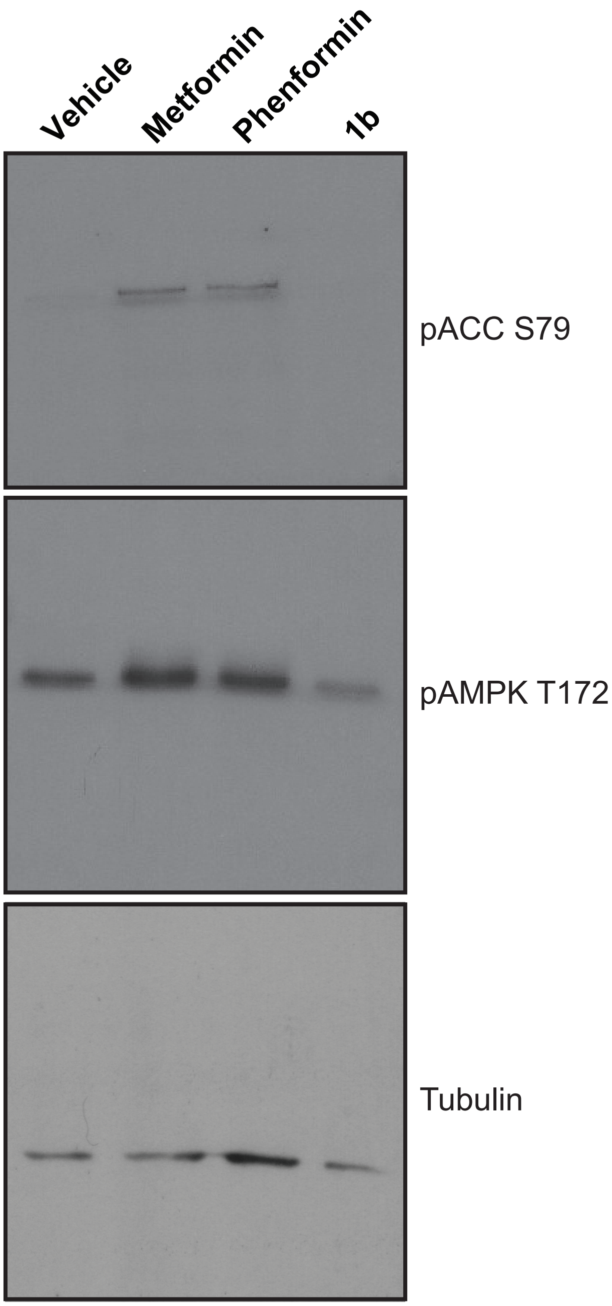
Vehicle
Metformin
Phenformin
1b
pACC S79
pAMPK T172
Tubulin

1. Chen, W. W.; Freinkman, E.; Wang, T.; Birsoy, K.; Sabatini, D. M., Absolute Quantification of Matrix Metabolites Reveals the Dynamics of Mitochondrial Metabolism. *Cell* **2016,** *166* (5), 1324-1337 e11.
